# Cholinergic control of striatal GABAergic microcircuits

**DOI:** 10.1101/2021.11.22.469580

**Authors:** S. Kocaturk, F. Shah, B. Guven, J.M. Tepper, M. Assous

## Abstract

Cholinergic interneurons (CINs) are essential elements of striatal circuits and behaviors. While acetylcholine signaling via muscarinic receptors (mAChRs) have been well studied, more recent data indicate that postsynaptic nicotinic receptors (nAChRs) located on GABAergic interneurons (GINs) are equally critical. One demonstration is that CINs stimulation induces large disynaptic inhibition of SPNs mediated by nAChR activation of striatal GINs. While these circuits are ideally positioned to modulate striatal output activity, the neurons involved are not definitively identified due largely to an incomplete mapping of CINs-GINs interconnections. Here, we show that CINs optogenetic activation evokes an intricate dual mechanism involving co-activation of pre- and postsynaptic mAChRs and nAChRs on four GINs populations. Using multi-optogenetics, we demonstrate the participation of tyrosine hydroxylase-expressing GINs in the disynaptic inhibition of SPNs likely via heterotypic electrical coupling with neurogliaform interneurons. Altogether, our results highlight the importance of CINs in regulating GINs microcircuits via complex synaptic/heterosynaptic mechanisms.

## Introduction

The striatum possesses the highest density of acetylcholine (ACh) in the basal ganglia. Striatal cholinergic interneurons (CINs) are responsible for the bulk of striatal ACh although extrinsic cholinergic afferents from brainstem structures may also play a role ^8,9^.

The effect of ACh in the striatum through muscarinic receptors (mAChRs) have received particular attention in part due to their wide cellular distribution. Indeed, mAChRs are expressed on the axon terminals of most striatal afferents as well as by all striatal neurons examined including the SPNs (for review see ^11,12^). However, the functional role of mAChRs expressed by striatal GABAergic interneurons (GINs) is less known. The few electrophysiological studies report that presynaptic mAChRs negatively regulate synaptic connections between fast-spiking interneurons (FSIs) and spiny projection neurons (SPNs) ^13^ and activation of postsynaptic mAChRs in low-threshold spike interneurons (LTSIs) has a hyperpolarizing effect ^6,14^.

The role of nicotinic receptors (nAChRs) in striatal circuits has until recently been relatively understudied^12^. The main role of nAChRs in the central nervous system had been suggested to be the modulation of the release of neurotransmitters via presynaptic receptors ^15–17^. Perhaps the most studied role of nAChRs in this regard involves their expression on presynaptic dopaminergic or glutamatergic afferents ^18–20^. However, we now know that CINs do not solely operate either as neuromodulatory neurons monosynaptically innervating SPNs via mAChR or through the modulation of neurotransmitter release. Rather, they can also exert fast nicotinic synaptic effects in the network by activating subpopulations of striatal nAChR-expressing GINs as has been observed in other brain structures including the cortex or hippocampus ^21–23^.

This notion was first suggested by indirect evidence demonstrating that nAChRs activation depolarizes striatal FSIs and facilitates GABA-mediated inhibition of the SPNs ^13,24,25^. Subsequently, we demonstrated that optogenetic activation of CINs elicits very large, disynaptic compound GABAergic IPSP/Cs on SPNs that are secondary to β2-subunit containing nAChR (β2-nAChR) activation of some populations of striatal GINs ^2–4,26^. The optically elicited IPSCs are kinetically biphasic consisting of a fast and a slow component (fIPSC and sIPSC), respectively mediated by GABA_Afast_ and GABA_Aslow_ currents ^2^. While this nicotinic-mediated circuit is positioned to play a central role in the regulation of striatal neuronal activity and related behaviors, its component composition is still mostly unknown. While the sIPSC (GABA_Aslow_) seems to rely, at least in part, on nicotinic activation of neurogliaform interneurons (NGFs), the source(s) of the fIPSCs (GABA_Afast_) is still unclear but seems to involve multiple GINs targeted in the Htr3a-Cre mice ^2,3^ (but see ^4^). The incomplete description of these circuits and their functional roles care due to the lack of a thorough mapping of the interactions between CINs and striatal GINs.

Here, we performed a comprehensive circuit analysis study of the interconnections between CINs and the different populations of striatal GINs using optogenetic stimulation in double transgenic mice associated with detailed pharmacology. Further, combining excitatory and inhibitory opsins, we were able to dissect the participation of distinct populations of striatal GINs in the disynaptic inhibition of SPNs. Our results demonstrate a heretofore unknown organizational complexity of the CINs-GABAergic interneurons circuits which, via their participation in the nicotinic-mediated circuit described above, play an important role in the regulation of striatal output activity. This study also underlies the relevance of understudied postsynaptic nAChRs in the striatum and their potential implications for the development of targeted pharmacological approaches to treat associated disorders of the striatum and basal ganglia such as Parkinson’s disease, dystonia, Tourette syndrome or nicotine addiction.

## Results

### CIN innervation of THINs

Our general approach to identify the innervation of different populations of striatal GINs by CINs was to generate double transgenic mice in which CINs natively express ChR2-eYFP (ChAT-ChR2-EYFP, Suppl. Fig 1) crossed with different Cre-expressing lines targeting different populations of GINs. The injection of a Cre-dependent tdTomato (AAV5-Flex-tdTomato) in the striatum permits selective fluorescent visualization of the targeted GIN population. Using this approach, we have previously demonstrated the cholinergic innervation of NGF and FAIs as well as the pharmacology involved ^2,10^ (Suppl. Fig. 2)

**Figure 1.**
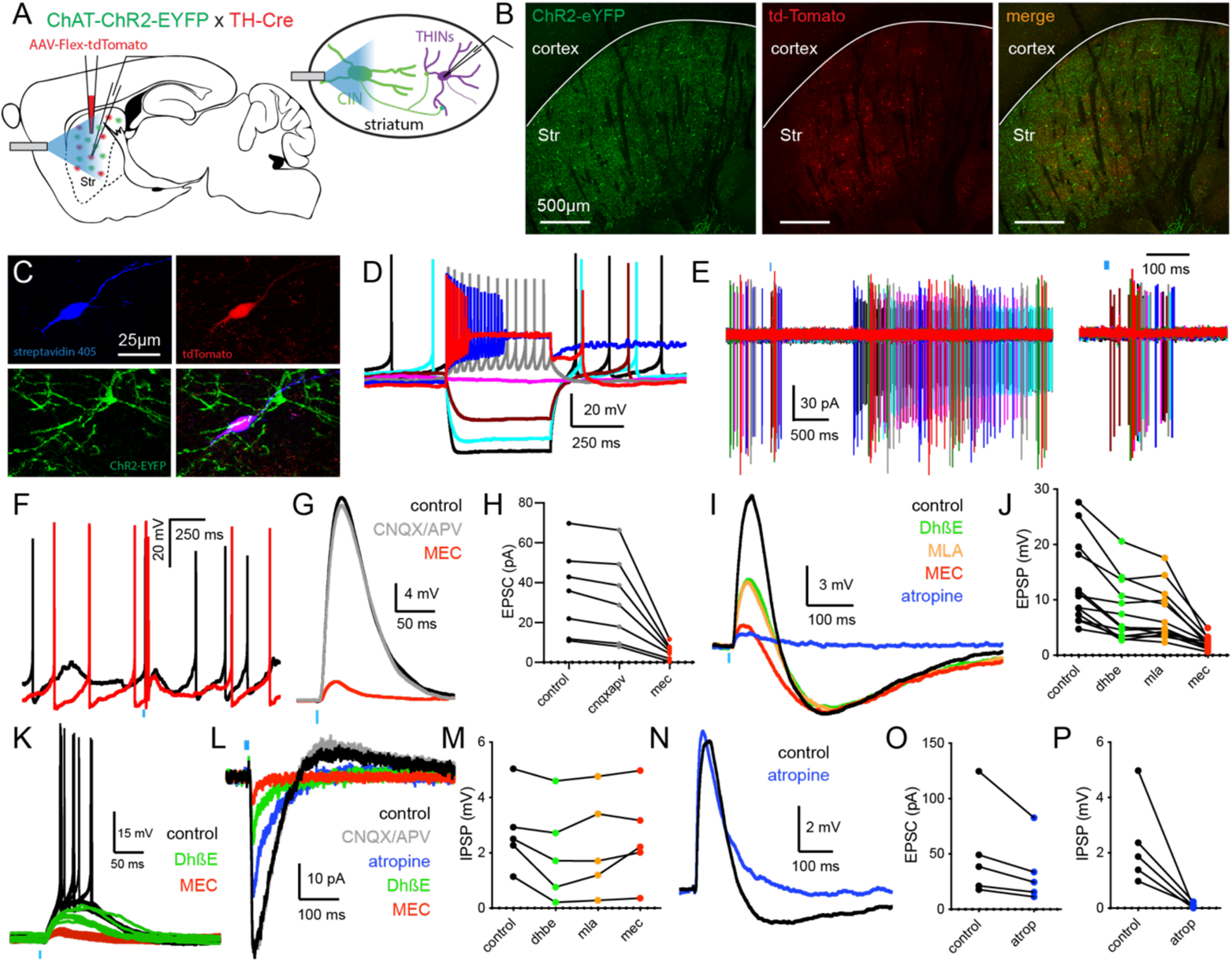
Cholinergic innervation of THINs. **A**. Schematic illustrating the experimental design using double transgenic ChAT-ChR2-eYFP::TH-Cre mice injected with a Cre dependent tdTomato AAV in the striatum. **B**. Confocal pictures showing striatal transduction of THINs (red, tdTomato) and CINs (green, ChR2-eYFP). **C**. Confocal pictures of a recorded THIN expressing tdTomato, filled with biocytin (revealed with streptavidin 405) and surrounded by ChR2 axons (green). **D**. Typical response of a THIN to negative and positive somatic current injection. **E**. Cell-attached recording of a THIN which respond to optogenetic stimulation of CINs (blue bar) with a burst of APs. Right: Enlargement around the stimulation perod. **F**. Current clamp recording of a THINs firing APs in response to CINs stimulation. **G**. The CINs-induced EPSP is slightly reduced by glutamate receptor antagonists (grey, CNQX, 10μM and APV, 10μM, blockers of AMPA and NMDA receptors, respectively) and dramatically affected by subsequent application of a nAChR antagonist (mecamylamine, 5μM, MEC, quantified in **H**). **I**. Current clamp recording and **J**. Quantification of the effect of selective ACh receptor antagonist on the CIN-induced EPSP. **K**. Individual current clamp traces showing that CINs stimulation induced AP firing in a THIN (black traces). Bath application of DhβE reduces the EPSP and prevent AP firing (green traces). Subsequent application of MEC almost completely abolish the EPSP (red traces). **L**. Voltage clamp recording representing the response of a THINs to optogenetic stimulation of CINs. The EPSC is often followed by an IPSC. While the EPSC is mostly affected by nAChRs antagonists (DHβΕ, 1μM and MEC, 5μΜ) the IPSC is abolished by a mAChR antagonist (Atropine, 10μM). **M**. Quantification of the effect of nAChRs antagonists on the IPSP. **N**. Current clamp recording showing the effect of atropine on the IPSP (quantified in **P**). **O**. Quantification of the effect of atropine on the CINs-induced EPSC.

**Figure 2.**
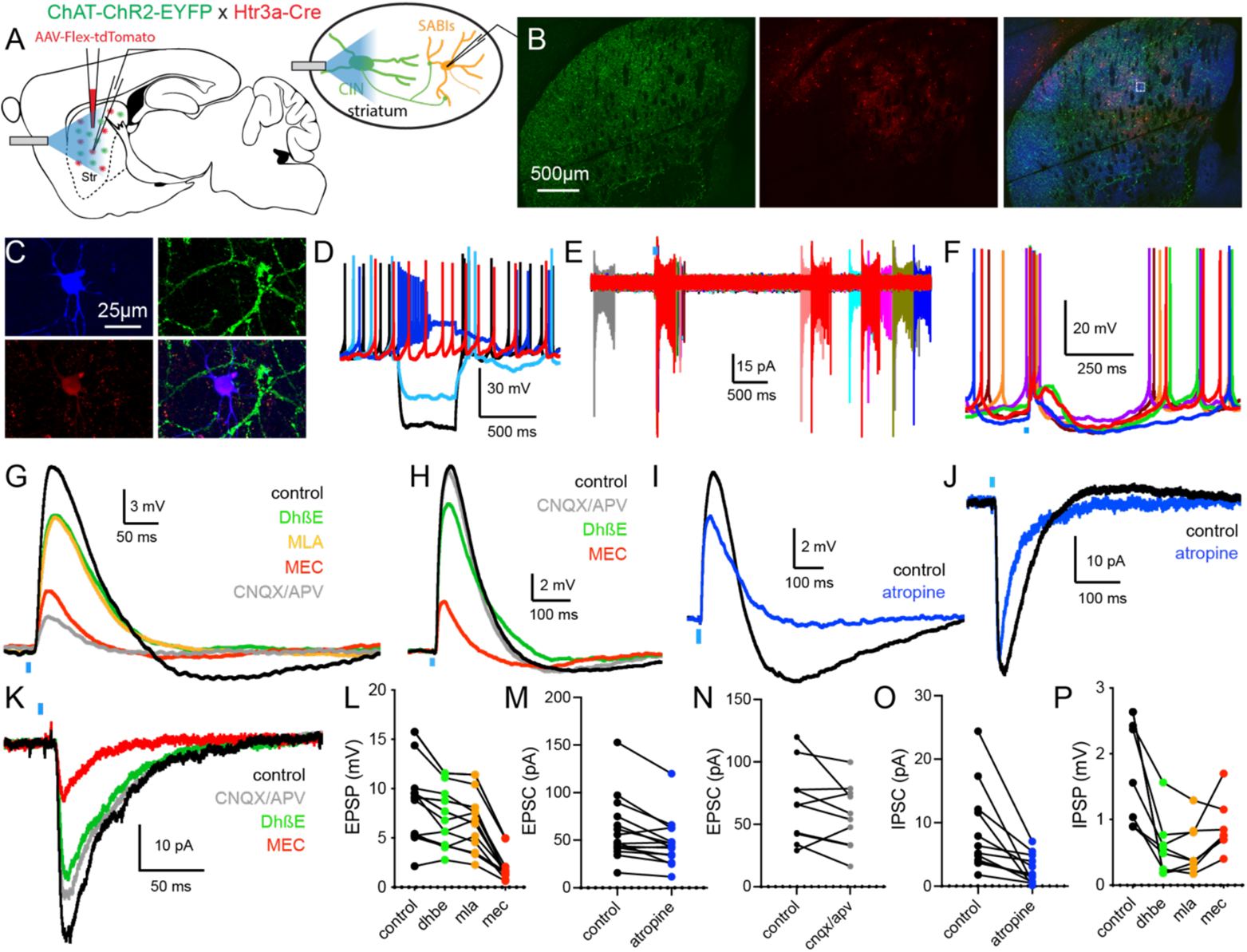
Cholinergic innervation of SABIs. **A**. Schematic illustrating the experimental design using double transgenic ChAT-ChR2-eYFP::Htr3a-Cre mice injected with a Cre dependent tdTomato AAV in the striatum. **B**. Confocal pictures showing striatal transduction of Htr3a-Cre interneurons (red, tdTomato) and CINs (green, ChR2-eYFP). **C**. Confocal pictures of a recorded SABI expressing tdTomato, filled with biocytin (streptavidin 405) and surrounded by ChR2 axons (green). **D**. Typical response of a SABI to negative and positive somatic current injection. **E**. Typical cell-attached recording of a SABI showing spontaneous burst activity. The SABI respond to optogenetic stimulation of CINs (blue bar) with a large burst of APs. **F**. Current clamp recording of a SABI firing APs in response to CINs stimulation. **G,H,I,J,K**. The CINs-induced EPSP/C is not significantly affected by bath application of glutamate receptor antagonists (grey, CNQX, 10μM and APV, 10μM, blockers of AMPA and NMDA receptors, respectively, quantified in **N**). However, we measured a significant reduction of the EPSP/C after application of DHβE (1μM, green) and MEC (5μM, red) by not by MLA (500nM, orange, quantified in **L**). Further, the mAChR antagonist atropine significantly reduce the EPSP/C amplitude (10μΜ, blue, quantified in **M**). In SABIs, the CIN-induced EPSP/C is often followed by an IPSP/C (**G,I,J**) which can be significantly reduced by DHβE (quantified in **P**) and completely abolished by atropine (quantified in **O**).

To examine the connectivity between CINs and THINs, we used double transgenic ChAT-ChR2-eYFP::TH-Cre mice and transduced THINs with td-Tomato as described above (Fig 1 A-C). CINs were spontaneously active, presented characteristic intrinsic electrophysiological properties as previously described and responded to brief blue light pulses (2 ms) with action potential (AP) firing consistent with ChR2 expression (Suppl. Fig 1).

Similarly, THINs presented characteristic intrinsic electrophysiological properties of the type I THINs ^7,27,28^ being spontaneously active both in whole-cell and cell-attached mode, a high input resistance and the presence of plateau potential following depolarizing current injection (Fig 1 D,E). Optogenetic activation of CINs (2 ms blue light pulse) evoked large EPSP/Cs in almost all recorded THINs (n = 27/28, EPSP amplitude: 11.10 ± 1.43 mV, τ=73.48 ± 3.97 ms, EPSC amplitude Vh −70mV: −42.64 ± 7.73 pA Fig. 1 F,G,I,K,L,N) that were sufficient to trigger AP firing in the majority of them (1-3 spikes; 74.07%; n=20/27; Figure 1 E-G, K). In most THINs the cholinergic-induced EPSP/C was followed by an IPSP/C exhibiting slower kinetics (n=23/27, IPSP amplitude: −1.57 ± 0.27 mV, τ= 725.34 ± 158.3 ms, Fig. 1 E,I,L,N), able to induce a pause in the firing activity of THINs.

First, we tested the effect of glutamate receptor antagonists on the EPSC amplitude. Bath application of AMPA and NMDA receptor antagonists (respectively CNQX, 10 μM and APV, 10μM) reduced slightly but significantly the amplitude of the cholinergic-mediated excitatory response (−3.68 ± 0.67 pA, [−10.56%] p=0.0016, t= 5.472, two-tailed paired t-test, n=7, Fig. 1 H). Further application of a nicotinic receptor antagonist largely abolished the EPSC (−25.63 ± 6.81 pA vs CNQX/APV condition, [−82.2%] p=0.0094, t= 3.761, two-tailed paired t-test, n=7, Fig. 1 G-L). Then we tested the subtype of nAChR involved in the EPSP/C by using selective nAChR antagonists. Bath application of dihydro-β-erythroidine (DhβE, a type II antagonist for nAChR containing β2* subunits, 1 µM) significantly reduced the amplitude of the excitatory response (−4.82 ± 0.87 mV, [−37.81%] vs. control; Fig 1 I-L). The remaining EPSP/C was not affected by methyllycaconitine (MLA, an α7* containing nAChR antagonist, 500 nM) but was almost completely abolished by mecamylamine (MEC, 5 µM, −5.76 ± 1.33 mV [−72.6%] vs DhβE; one-way ANOVA, F (1.354, 16.25) = 25.43, Tukey’s multiple comparisons: control vs. DhβE: p=0.0006; DhβE vs. MLA: p=0.42; MLA vs. MEC: p=0.0046, n=13, Fig. 1 I-L) indicating a mixed type II and III nicotinic receptor composition in THINs. In additional pharmacological experiments we tested the involvement of muscarinic receptors in the EPSP/Cs induced by the optogenetic stimulation of CINs. Application of atropine (10 μM) did not significantly reduce the amplitude of the EPSP/C (p=0.065, n=5, Fig. 1 N,O). However, while nAChR antagonists did not have a significant effect on the IPSP/C (DhβΕ: −0.77 ± 0.22 mV vs control; MLA: +0.27 ± 0.13 mV vs. DhβE; MEC: +0.28 ± 0.21 mV vs. MLA; one-way ANOVA, F (1.853, 7.411) = 4.97, Tukey’s multiple comparisons: control vs. DhβE: p=0.0819; DhβE vs. MLA: p=0.29; MLA vs. MEC: p=0.59, n=5), atropine incubation almost completely abolished the IPSP/Cs (−2.22 ± 0.71 mV [−95.7%] vs. control, p=0.035, t= 3.12, two-tailed paired t-test, n=5, Fig. 1 M-P). Further, using paired recording of CINs and THINs we confirmed the direct synaptic connection between these cells (^1^ Suppl. Fig 3).

**Figure 3.**
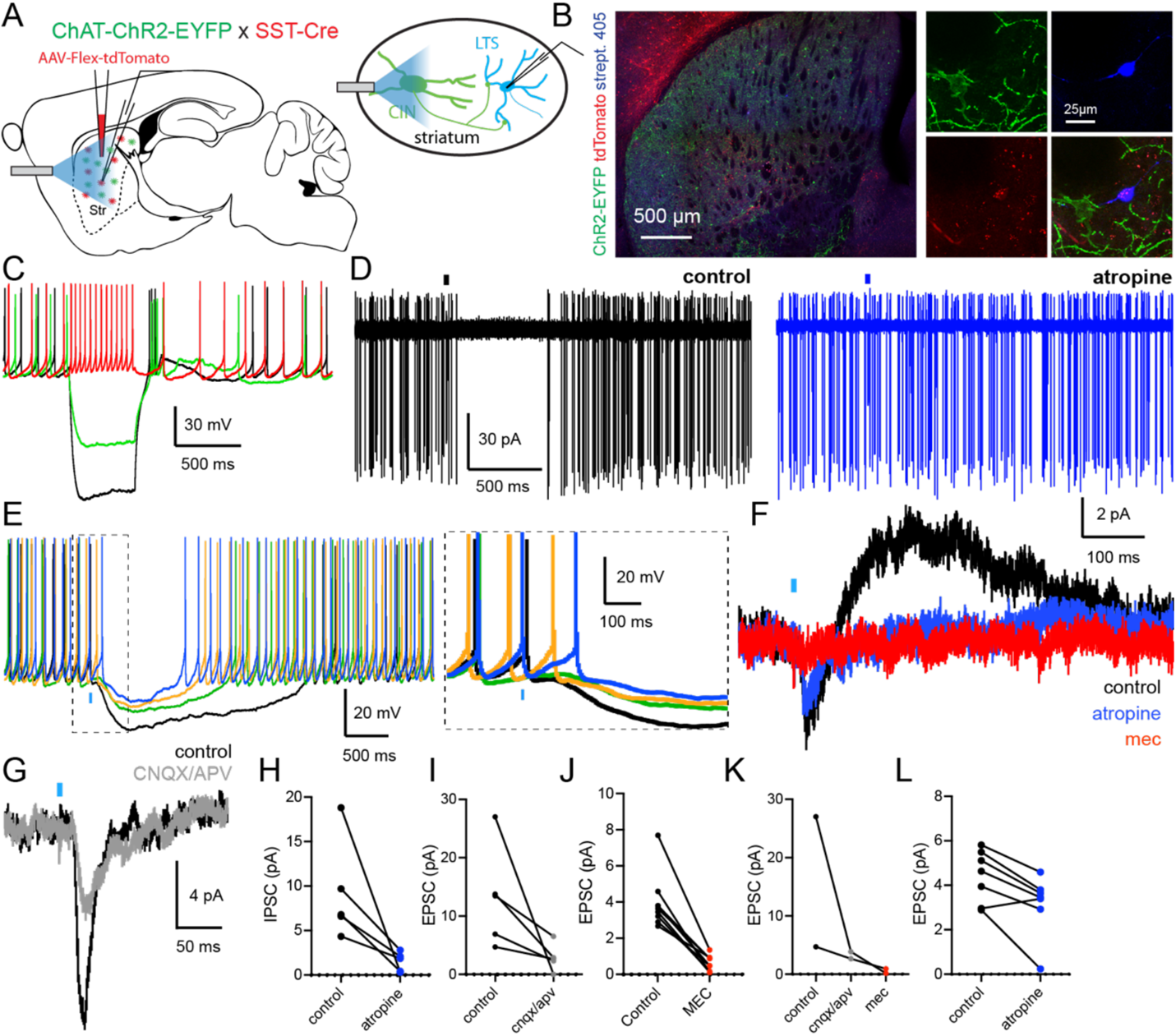
Cholinergic innervation of LTSIs. **A**. Schematic illustrating the experimental design using double transgenic ChAT-ChR2-eYFP::SST-Cre mice injected with a Cre dependent tdTomato AAV in the striatum. **B**. Left: Confocal pictures showing striatal transduction of LTSIs (red, tdTomato) and CINs (green, ChR2-eYFP). Right: Confocal pictures of a recorded LTSI expressing tdTomato, filled with biocytin (streptavidin 405) and surrounded by ChR2-expressing axons/cells (green). **C**. Typical response of a LTSI to negative and positive somatic current injection. **D**. Cell-attached recording of a LTSI showing spontaneous tonic activity. LTSI respond to optogenetic stimulation of CINs (blue bar) with a few APs followed by a long pause. **E**. Current clamp recording of a spontaneously active LTSI responding to CINs stimulation with an AP followed by a long pause. Right: enlargement of the dashed squared window. **F**. Voltage clamp recording. CINs stimulation evoke a modest EPSC followed by a slow IPSC (black trace). Application of atropine (10μM, blue trace) mostly affect the IPSC (quantified in **H,L**). Application of MEC (5μM, red trace) abolishes the EPSC (Quantified in **J, K**). **G**. Application of glutamate receptor antagonists (grey, CNQX, 10μM and APV, 10μM) reduces the EPSC amplitude (quantified in **I,K**).

These results demonstrate that CINs innervate striatal THINs with an intricate dual effect involving complex receptor pharmacology with type II and type III nAChR-mediated excitation and a muscarinic-mediated slow kinetic inhibition.

### CIN innervation of SABIs

To examine the connectivity between CINs and SABIs, we used double transgenic ChAT-ChR2-eYFP::Htr3a-Cre mice and transduced Htr3a interneurons with td-Tomato as described above (Fig. 2 A-C). As we described in previous studies ^3,7,10^, targeted GINs populations in the Htr3a-Cre mice include FSIs, NGFs as well as FAIs and SABIs. SABIs were readily identifiable based on their characteristic spontaneous bursty activity pattern, very high input resistance, depolarization block after injecting positive somatic current injection and the lack of connectivity with SPNs (^7,29^, Fig. 2 D-E, Suppl. Fig. 4). Optogenetic activation of CINs (2 ms blue light pulse) evoked large EPSP/Cs in all recorded SABIs (n = 67, EPSP amplitude: 11.49 ± 0.85 mV, τ=93 ± 9.91 ms, n= 57; EPSC amplitude Vh −70mV: −65.51 ± 6.34 pA, τ=49.8 ± 16.7 ms, n=48, Fig. 2 F-K) that were sufficient to trigger AP firing and characteristic large bursts when recorded in cell attached (Fig. 2 E,F). In most SABIs, the cholinergic-induced EPSP/C was followed by an IPSP/C presenting a slower kinetic (n=54/67, IPSP amplitude: −2.38 ± 0.27 mV, n=41, τ= 412.8 ± 64.02 ms, IPSC amplitude: 7.88 ± 1.07 pA, n=32, τ= 278.7 ± 31.8 ms, Fig. 2 G-K).

**Figure 4.**
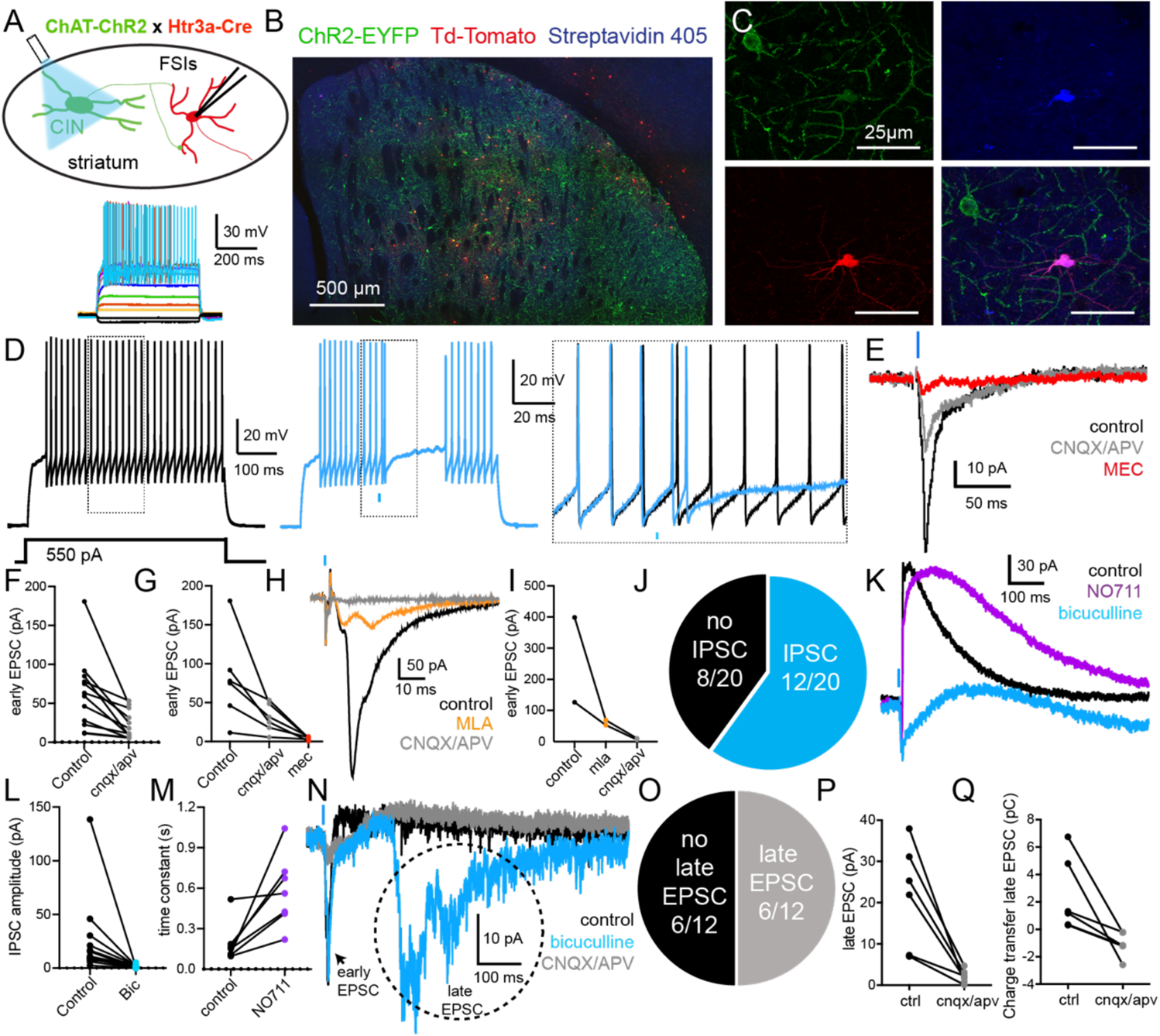
Cholinergic innervation of FSIs. **A**. Schematic illustrating the experimental design using double transgenic ChAT-ChR2-eYFP::Htr3a-Cre mice injected with a Cre dependent tdTomato AAV in the striatum. Bottom: Typical response of a FSI to negative and positive somatic current injection. **B**. Confocal pictures showing striatal transduction of Htr3a-Cre interneurons (red, tdTomato) and CINs (green, ChR2-eYFP). **C**. Confocal pictures of a recorded FSI expressing tdTomato, filled with biocytin (streptavidin 405) and surrounded by ChR2-expressing axons/cells (green). **D**. Left (black): Large somatic current injection triggers AP firing in a recorded FSI. Middle (blue): Optogenetic stimulation of CINs evokes a short increase in AP firing followed by a pause in current-induced firing activity. Right: Enlargement around the optogenetic stimulation. **E**. Voltage clamp recording showing a CIN-induced EPSC in a FSI (black). The EPSC is significantly reduced by bath application of glutamate receptor antagonists (grey, CNQX, 10μM and APV, 10μM) and almost completely abolished by subsequent addition of the nAChR antagonist, MEC (5μΜ, red, quantified in **F,G**). **H**. The CIN induced EPSC in FSI is significantly reduced by bath application of an α7-nAChR antagonist (methyllycaconitine, MLA, 500nM, quantified in **I**). **J-N**. When voltage clamped at −45mV, the majority of FSI respond the CINs optogenetic stimulation with an IPSC following the EPSC (n=12/20, **J**). **K**. The kinetic of the IPSC is significantly increased by application of the GABA transport blocker (purple, NO711, 10μM, quantified in **M**). Further, the IPSC is completely abolished by a GABA_A_ receptor antagonist (blue, bicuculline, 10μM, quantified in **L**). **N-P**. Application of bicuculline revealed a late excitatory phase occurring several 100s of ms following CINs optogenetic stimulation. This was observed in n=6/12 FSIs recorded in these conditions (Vh=−45mV, bicuculline 10μM, **N,O**). These EPSC barrages can be blocked by bath application of glutamate receptor antagonists (grey, CNQX, 10μM and APV, 10μM, quantified in **P**: amplitude and **Q**: charge transfer).

First, we tested the effect of glutamate receptor antagonists on the EPSC amplitude. Bath application of AMPA and NMDA receptor antagonists (respectively CNQX, 10 μM and APV, 10μM) did not affect the amplitude of the excitatory response following CINs optogenetic activation (−9.35 ± 5.66 pA, [−14.1%] p=0.133, t= 1.652, two-tailed paired t-test, n=10, Fig. 2 H,K,N), suggesting that EPSP/Cs was mediated principally by activation of nAChRs. Application of DhβE (1 µM) slightly but significantly reduced the amplitude of the excitatory response (−1.64 ± 0.41 mV, [−18.3%] vs. control; Fig. 2 G,H,K,L). The remaining EPSP/C was not affected by MLA (500 nM) but was greatly reduced by mecamylamine (MEC, 5 µM, −4.78 ± 0.46 mV [−69.4%] vs MLA; one-way ANOVA, F (1.558, 21.81) = 77.4, Tukey’s multiple comparisons: control vs. DhβE: p=0.0067; DhβE vs. MLA: p=0.248; MLA vs. MEC: p=3*10^−7^, n=15) indicating a mixed type II and III nicotinic receptor composition in SABIs. In additional pharmacological experiments we tested the involvement of muscarinic receptors in the EPSP/Cs induced by the optogenetic stimulation of CINs. Application of atropine (10 μM) also significantly reduced the amplitude of the EPSP/C (−12.9 ± 3.48 pA, [−21.3%], p=0.0023, t=3.71, n=15, two-tailed paired t-test, Fig 2 I,J,M). Interestingly, the IPSC/P was significantly reduced by bath application of atropine (−6 ± 1.72 pA, [−69.77%], p=0.005, t=3.496, n=12, two-tailed paired t-test) or DhβE but was not further decreased by subsequent application of MLA or MEC (DhβΕ: −1.23 ± 0.3 mV, [−68.05%] vs. control; MLA: +0.02 ± 0.06 mV vs. DhβE; MEC: −0.32 ± 0.11 mV vs. MLA; one-way ANOVA, F (1.340, 9.379) = 16.35, Tukey’s multiple comparisons: control vs. DhβE: p=0.018; DhβE vs. MLA: p=0.98; MLA vs. MEC: p=0.086, n=8, Fig 2 O,P).

These results demonstrate that CINs innervate striatal SABIs with a complex receptor pharmacology involving type II and type III nAChR-mediated excitation and a muscarinic-mediated slow inhibition.

### CIN innervation of LTSIs

Striatal LTSIs co-express neuropeptide Y (NPY)/nitric oxide synthase (NOS) and somatostatin (SST) ^30–32^. Hence, to examine the connectivity between CINs and LTSIs, we used double transgenic ChAT-ChR2-eYFP::SST-Cre mice and transduced LTSIs with td-Tomato as described above (Fig. 3 A-B). As we described in previous studies, LTSIs present characteristic intrinsic electrophysiological properties with spontaneous AP firing (both in cell-attached mode and whole-cell recordings), high input resistance and low threshold Ca^2+^spikes (Fig. 3 C) ^6,30,33^. Optogenetic stimulation of CINs evokes a long (>500ms) pause response in spontaneously active LTSIs both in cell-attached as well as in whole-cell modes (Fig 3 D-E, ^6^). In some instances (25% of recorded LTSIs), the pause is followed by periods of bursts activity as previously described (Suppl. Fig 5, ^6^). However, we observed that the optogenetic-induced pause is preceded by an EPSP/C (7.12 ± 1.19 pA, τ=38.18 ± 5.25 ms, n= 25) sometimes sufficient to elicit AP firing in depolarized, spontaneously active LTSIs (Fig. 3 E-G). We dissected pharmacologically this EPSC by performing bath application of glutamatergic, nicotinic and muscarinic receptor antagonists. AMPA and NMDA receptor antagonists significantly reduced the EPSC (−10.3 ± 4.43 pA, [−78.2%], p=0.04, paired t-test, t=2.327, n=5, Fig. 3 G,I,K). Interestingly, a similar level of reduction of the EPSC amplitude was achieved by bath application of the nAChR antagonist (MEC, 5μM, −3.37 ± 0.45 pA, [−84.9%], two-tailed paired t-test, t=7.56, n=9, p=6.54 x 10^−5^, Fig. 3 F,J,K). In a few examples, we tried to test the impact of MEC on the remaining EPSC measured after glutamate receptor antagonist application. In a few LTSIs, subsequent application of MEC further reduced or even abolished the leftover EPSC, suggesting a mixed mechanism combining direct nAChR-mediated activation of LTSIs of moderate amplitude and a larger effect mediated by nicotinic-induced glutamate release from cortical and/or thalamic afferents (Fig. 3 K).

**Figure 5.**
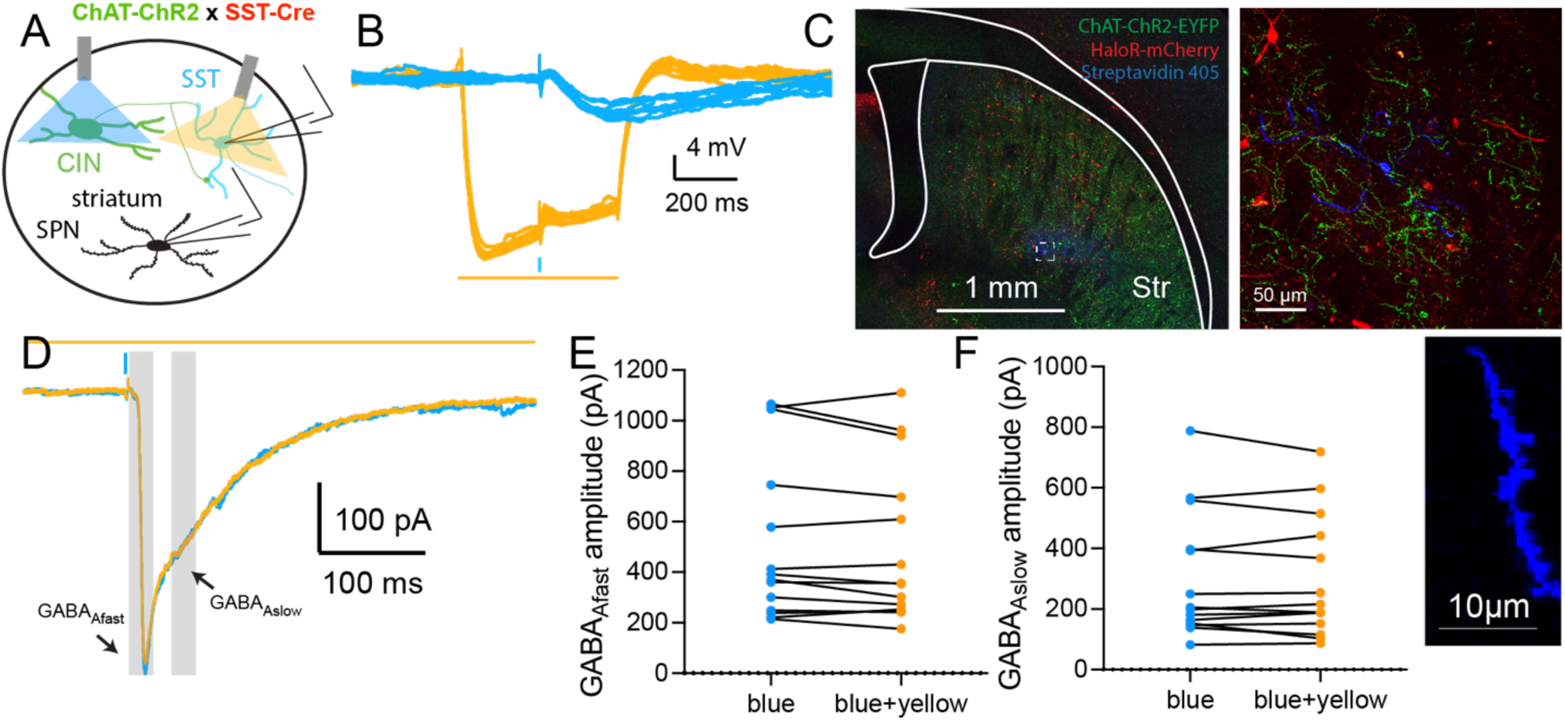
Influence of LTSIs in the nicotinic mediated disynaptic inhibition of SPNs. **A**. Schematic illustrating the experimental design using double transgenic ChAT-ChR2-eYFP::SST-Cre mice injected with a Cre dependent HR3.0 AAV in the striatum. This allows to optogenetically activate CINs with blue light and inhibit LTSIs with yellow light. **B**. Optogenetic activation of CINs with blue light evokes a small EPSP followed by an IPSP in an LTSI (blue traces). Concomitant inhibition of LTSIs with yellow light evokes a large hyperpolarization during the yellow light puse (500 ms) due to HR3.0 expression. **C**. Confocal pictures showing striatal transduction of LTSIs (red, HR3.0-mCherry) and CINs (green, ChR2-eYFP). SPNs were recoded with biocytin and revealed with Streptavidin 405nm. Bottom: Enlargement of a SPN dendrite showing abundant spines. **D**. Optogenetic activation of CINs evoke large disynaptic compound IPSC in SPNs (recorded with CsCl Internal, see methods) including a fast and a slow IPSC (GABA_Afast_ and GABA_Aslow_, respectively). Concomitant inhibition of LTSIs does not affect the amplitude of the GABA_Afast_ IPSC (**E**) and the GABA_Aslow_ IPSC (**F**).

Further, consistent with a previous study, we were able to block the cholinergic-induced pause in LTSIs with a muscarinic receptor antagonist (atropine, 10μM, ^6^). In voltage-clamp experiments, the pause exhibits slow kinetics (IPSC amplitude: 6.95 ± 0.77 pA, τ=390.3 ± 265.2 ms, n= 28, Fig. 3 D-F), similar to the IPSC observed in THINs and SABIs (see above). This slow IPSC was blocked by bath application of atropine (−7.75 ± 2.8 pA, [−83.86%] vs. control, p=0.0252, two-tailed paired t-test, t=2.77, n=5). Given that THINs innervate LTSIs ^5^ and CINs provide suprathreshold EPSP in THINs, we examined whether optogenetic stimulation of CINs could also evoke fast GABA_A_ IPSCs through this disynaptic circuit. When recording in voltage clamp at a holding voltage of −45mV, we measured the presence of polysynaptic fast IPSCs in n=4/8 tested LTSIs (Suppl. Fig. 5). The IPSCs were blocked by the GABA_A_ receptor antagonist, bicuculline (10 μM), demonstrating the existence of the disynaptic microcircuit.

### CIN innervation of FSIs

Next, we examined the responses of FSIs to optogenetic stimulation of CINs. We used double transgenic ChAT-ChR2-eYFP::Htr3a-Cre mice and transduced Htr3a interneurons with td-Tomato as described above (Fig. 4 A-C). As mentioned above striatal FSIs are among the GINs targeted in the Htr3a-Cre mice and represent a large proportion of the AAV-transduced cells ^10^. These FSIs exhibit all the characteristic intrinsic electrophysiological properties of previously identified parvalbumin (PV)-expressing striatal FSIs (hyperpolarized membrane potential, small input resistance and high frequency discharge, short duration AP firing, sometimes in interrupted bursts response to depolarizing current injection, Fig. 4 A) previously reported in rodents ^10,25,31,32^. Optogenetic stimulation of CINs evoked mixed excitatory/inhibitory response in FSIs revealed in current clamp recordings after eliciting AP firing by somatic current injection. Large somatic positive current injection in FSI elicited high frequency AP firing that can be interrupted for 10s of ms by optogenetic stimulation of CINs (Fig 4 D). This biphasic response of FSIs was investigated in voltage clamp recordings (Vh=−45mV). A single 2 ms optical pulse elicited an EPSC (−44.24 ± 11.09 pA, n=16, Fig. 4 E) possessing a fast decay time constant (τ=5.44 ± 0.65 ms) consistent with the co-involvement of both glutamate (CNQX/APV 10μΜ :-40.51 ± 10.63 pA [−64.58%] vs control, p= 0.0029, t=3.81, two-tailed paired t-test, n=12, Fig. 4 E-G) and nicotinic cholinergic transmission (MEC: −26.20 ± 7.28 pA [−86.53%] vs CNQX/APV condition, p=0.016, t=3.6, two-tailed paired t-test, n=6, Fig. 4 E, G-I, ^34^). Interestingly, similar elimination of the EPSC could be achieved via consecutive application of MLA and CNQX/APV suggesting the specific involvement of α7-containing nAChRs (n=2, Fig. 4 H,I). Further, in most cases, the EPSC is followed by an IPSC (n=12/20 recorded FSIs, 24.78 ± 10.02 pA, Fig. 4 J-M) with slow kinetic (τ=139.8 ± 49.23 ms) and properties of GABA_Aslow_ ^2,35–37^. The GABA_Aslow_ IPSC is significantly reduced by bath application of bicuculline (10μM: −23.15 ± 10.08 pA [−93.42%], p=0.04, t=2.296, two-tailed paired t-test, n=13, Fig. 4 K,L) and the decay time constant is greatly increased by blocking GABA reuptake (NO711, 10μΜ, τ control: 190.7 ± 55.9 ms, τ NO711: 579.3 ± 100.7 ms, [303.8% increase], n=7, p=0.0141, t=3.423, two-tailed paired t-test, Fig. 4 K,M). Unexpectedly, bath application of bicuculline unraveled a late phase of EPSCs barrages in a proportion of FSIs (Fig. 4 N-Q). These asynchronous EPSCs (amplitude: 21.77 ± 5.14 pA, charge transfer: 2.447 ± 1.09 pC, n=6/12 FSIs) could be blocked by glutamate receptor antagonists (CNQX/APV 10μM, [−90.2%], p=0.0107, t=3.964, two-tailed paired t-test, Fig. 4 N,P,Q) suggesting the existence of tonic GABAergic inhibition of glutamatergic release from unidentified intrastriatal terminals that is triggered by synchronous ACh release from CINs.

### Involvement of THINs and LTSIs in the disynaptic inhibition of SPNs

Synchronous activation of CINs evokes a disynaptic inhibition of SPNs composed of distinct GABA_Afast_ and GABA_Aslow_ components ^2–4^. These compound IPSCs are disynaptic and are due to activation of β2 subunit-containing nAChRs located on many different striatal GINs. While the GABA_Aslow_ has been demonstrated to be due to activation of β2*-nAChRs located on NGF interneurons that monosynaptically innervate SPNs, the source of the GABA_Afast_ current is still unclear but almost certainly involve, at least partially, striatal GINs targeted in the Htr3a-Cre mouse ^2–4^.

To test the involvement of LTSIs in the disynaptic inhibition of SPNs following synchronous activation of CINs, we used double transgenic mice ChAT-ChR2-eYFP::SST-Cre mice and transduced LTSIs with Halorhodopsin 3.0-mCherry (HR3.0), to allow activation of ChR2 and HR3.0 individually or simultaneously (Fig. 5 A-C). This enabled us to optogenetically disconnect the CINs-GINs-SPNs cell circuit on a trial-by-trial basis ^3^. As previously demonstrated optogenetic stimulation of CINs with brief blue light pulses (2 ms) elicited large IPSC in recorded SPNs (Vh=−70 mV, CsCl internal, see methods) which could be blocked both by β2*-nAChRs and GABA_A_ receptor antagonists (respectively DhβE, 1μM and bicuculline, 10μM) confirming the disynaptic nature of these IPSCs (Suppl. Fig 6, ^2,3^).

**Figure 6.**
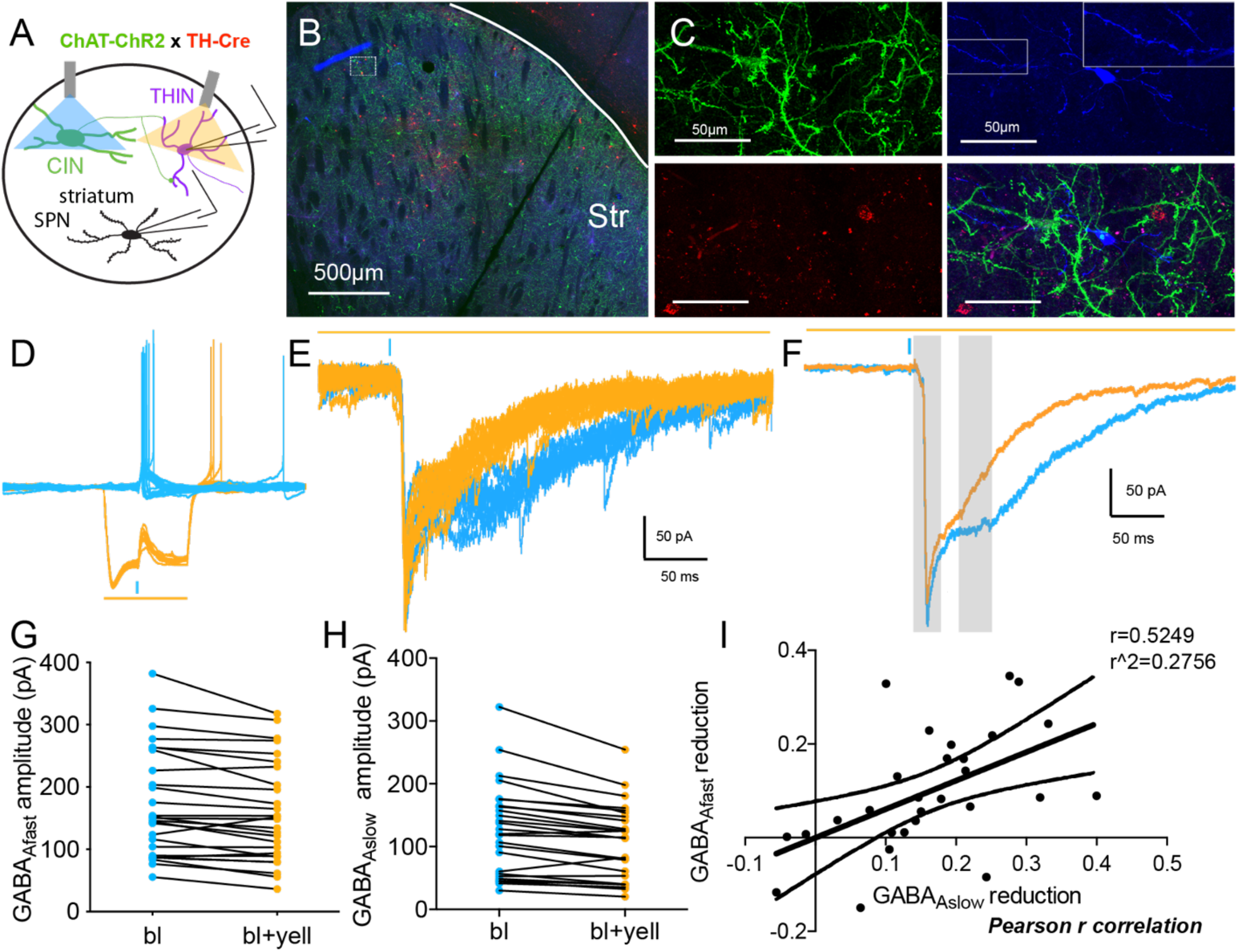
Influence of THINs in the nicotinic mediated disynaptic inhibition of SPNs. **A**. Schematic illustrating the experimental design using double transgenic ChAT-ChR2-eYFP::TH-Cre mice injected with a Cre dependent HR3.0 AAV in the striatum. This allows to optogenetically activate CINs with blue light and inhibit THINs with yellow light. **B-C**. Confocal pictures showing striatal transduction of THINs (red, HR3.0-mCherry) and CINs (green, ChR2-eYFP). **C**. SPNs were recoded with biocytin and revealed with Streptavidin 405nm. Inset: Enlargement of a SPN dendrite showing abundant spines. **D**. Optogenetic activation of CINs with blue light evokes large depolarization and AP firing in a THIN (blue traces). Concomitant inhibition of THINs with yellow light evokes a large hyperpolarization during the yellow light puse (500 ms) due to HR3.0 expression, preventing AP firing following stimulation of CINs. **E-F**. Optogenetic activation of CINs evoke large disynaptic compound IPSC in SPNs (recorded with CsCl Internal, see methods) including a fast and a slow IPSC (GABA_Afast_ and GABA_Aslow_, respectively, blue traces). Concomitant inhibition of THINs induced a reduction in the GABA_Afast_ and the GABA_Aslow_ (**E**: Examples of individual traces, **F**: average responses). Quantification of the reduction of the GABA_Afast_ (**G**) and the GABA_Aslow_ (**H**) following optogenetic inhibition of THINs. **I** Pearson r analysis demonstrating that the reduction of the GABA_Afast_ and the GABA_Aslow_ are correlated.

Stimulation of CINs (2 ms blue light pulse) evoked small amplitude depolarization in LTSIs, evoking AP firing on rare occasions. Consistent with these results, optogenetic inhibition of LTSIs with yellow light did not significantly reduce the amplitude of either GABAergic current in SPNs following CIN optogenetic stimulation (GABA_Afast_:-20.98 ± 13.29 pA, p=0.14; GABA_Aslow_: −6.37 ± 8.99 pA, p=0.49, two-tailed paired t-test, Fig. 5 D-F). These results show that LTSIs are not significantly involved in this nicotinic mediated circuit.

Then, we tested the involvement of THINs in the disynaptic inhibition of SPNs following stimulation of CINs using the same double optogenetic approach. In ChAT-ChR2-eYFP::TH-Cre mice (Fig. 6 A-C), optogenetic stimulation of CINs evoked large depolarization and AP firing in recorded THINs (Fig. 6 D) consistent with the results described above (Fig. 1). On alternate trials, we optogenetically inhibited THINs transduced with HR3.0 with yellow light (590nm, 700 ms, starting 200 ms before CINs stimulation). The yellow light significantly hyperpolarized THINs and blocked AP firing in response to CINs stimulation, leaving a subthreshold depolarization (n=7, Fig. 6 D). This confirms the efficacy of this double optogenetic approach demonstrating that blue light pulse evokes AP firing in THINs as previously described and that halorhodopsin-induced hyperpolarization is sufficient to prevent the CINs-induced AP firing in THINs.

Next, we repeated the same stimulation protocol while recording from SPNs in voltage clamp. Optogenetic activation of CINs evoked a large compound IPSC in SPNs consisting often of distinct GABA_Afast_ and GABA_Aslow_ components (GABA_Afast_ amplitude: 205.4 ± 17.41 pA, latency: 17.88 ± 0.45 ms; GABA_Aslow_ amplitude: 151.6 ± 14.39 pA, τ GABA_Aslow_: 73.05 ± 6.4 ms; n=20 SPNs). Simultaneous illumination with yellow light that selectively inhibited THINs, produced a relatively small but significant reduction in the amplitude of the GABA_Afast_ (−16.32 ± 4.64 pA [−7.95%], p=0.0023, t=3.514, two-tailed paired t-test), demonstrating that THINs, due to their striatal synaptic connectivity (receiving nicotinic activation from CINs and inhibiting SPNs via GABA_Afast_ ^7,27,28^), participate in this circuit (Fig. 6 E-I). More surprisingly, the inhibition of THINs produces a more important reduction of the GABA_Aslow_ (−22.37 ± 4.56 pA [−14.8%], p<0.0001, two-tailed paired t-test). We verified that this effect was not a light artefact by reproducing similar optogenetic stimulation protocols in single transgenic ChAT-ChR2-eYFP mice (Suppl. Fig. 6). This was unexpected as THINs do not monosynaptically innervate SPNs via GABA_Aslow_ ^7,27,28^. As mentioned above, the GABA_Aslow_ induced in SPNs was attributed to CIN activation of NGFs, activation of which evokes GABA_Aslow_ IPSC in SPNs ^2,30^. Given the ability of NGFs to form heterotypic gap junction with other interneurons in other structures such as the cortex and hippocampus ^38–41^, we hypothesized that striatal NGFs may be electrically coupled with THINs and that this mechanism would explain why optogenetic manipulation of THINs modulates GABA_Aslow_ in SPNs. ***Electrical coupling between THINs and NGFs***

One of the explanations for the reduction of the GABA_Aslow_ in SPNs following optogenetic inhibition of THINs would be that THINs and NGFs are electrically coupled through gap junctions (Fig. 7). To test this, we measured responses elicited in NGF interneurons by the optogenetic activation of THINs (Fig 7A-B). This was performed in double transgenic TH-Cre::NPY-GFP mice injected with a Cre-dependent ChR2 AAV in the striatum. Strikingly, we found that 55.6% of NGFs recorded (n=10/18) respond to THINs stimulation with an inward current (amplitude: 14.05 ± 2.82 pA, n=10, Fig. 7 C-L) that show little or no variation in amplitude when THINs were stimulated with a train (5 pulses, 20Hz, p1: 11.73 ± 1.75 pA, p2: 11.45 ± 1.45 pA, p3: 10.31 ± 1.28 pA, p4: 9.81 ± 1.23 pA, p5: 9.69 ± 1.3 pA, Fig. 7 F-H,J,K) and a relatively slow decay time constant (τdecay: 25.21 ± 3.5 ms, Fig. 7 L). All of these are consistent with the slow inward current resulting from electrical coupling between THINs and NGFs. When investigating further, optogenetic activation of THINs evoked spikelets in NGFs also consistent with the existence of electrical coupling (Fig. 7 C-D). Finally, application of carbenoxolone, 100μM, which disrupts gap junction coupling ^42^, caused a dramatic reduction of the amplitude of the slow inward current in NGF interneurons following optogenetic stimulation of THINs (−8.82 ± 0.49 pA [−67.7%], p=0.0031, two-tailed paired t-test, t=18.03, n=3, Fig. 7 H,K). Altogether these results demonstrate the existence of functional electrical coupling between THINs and NGFs. These also indicate that the decrease in the disynaptic GABA_Aslow_ measured in SPNs following optogenetic stimulation of CINs and inhibition of THINs is indirect due to NGFs hyperpolarization as a result of heterotypic electrical coupling.

**Figure 7.**
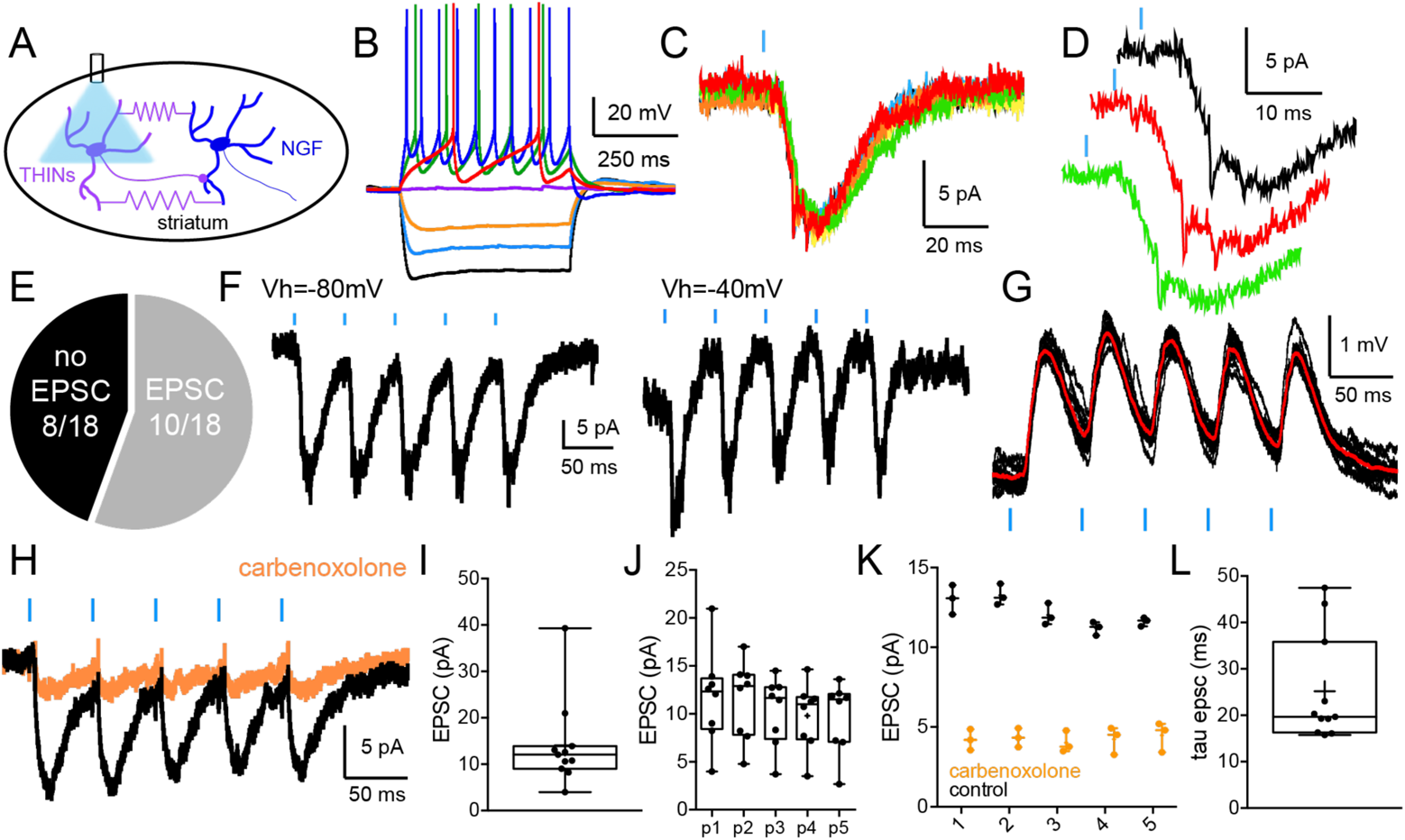
Electrical coupling between THINs and NGF interneurons. **A**. Schematic illustrating the experimental design using double transgenic TH-Cre::NPY-GFP mice injected with a Cre dependent ChR2 AAV in the striatum. This allows to optogenetically activate THINs with blue light record from NGF interneurons. **B.** Typical response of an NGF interneuron to somatic current injection. **C.** Optogenetic stimulation evokes inward current in n=10/18 recorded NGFs (**E**). **D**. The inward current often present “spikelets” indicative of electrical coupling. **F**. The inward current does not reverse at −40mV demonstrating that this is not a GABA_A_-mediated response. Further, in a train stimulation (5 pulses at 20Hz) the amplitude of the EPSP/C do not significantly change (**F,G,H** and quantified in **I** and **J**). Bath application of a gap junction blocker (carbenoxolone, 100μΜ) dramatically reduce the size of the inward current. **L**. Quantification of the decay time constant of the inward current.

## Discussion

Postsynaptic striatal nAChRs containing the β2* subunit, selectively expressed by striatal interneurons, are known to be involved in the powerful disynaptic inhibition of SPNs and play an important role in the regulation of striatal neuronal activity ^2–4^. These circuits provide a way for CINs to rapidly control striatal output. However, the multiple circuits and different interneurons involved have not previously been precisely understood. One reason has been the largely incomplete mapping and pharmacology of the interconnections between CINs and the increasing diversity of striatal GINs and their intrastriatal connectomes. Here, we demonstrate that CINs innervate at least 5 populations of GINs involving different synaptic circuits and nAChR subtypes. During this circuit analysis, we were able to test, using double optogenetics, the participation of the different populations of GINs in the disynaptic inhibition of SPNs induced by CINs population stimulation.

It has been shown that THINs express functional nAChRs as they respond strongly to local application of agonists ^43,44^. Further, using multiple simultaneous whole cell recordings, it has been demonstrated that CINs directly innervate THINs ^1^. Here, we examined the response of THINs to the optogenetic stimulation of CINs as well as the pharmacology involved. Our results demonstrate an atypical dual control of THINs activity by CINs. Optogenetic stimulation of CINs evokes a large suprathreshold nicotinic-mediated excitatory responses in THINs involving partially β2*-nAChR as blockade of these receptors prevents cholinergic-induced AP firing in THINs. The residual part of the EPSP/C involve a type of nAChR distinct from the α7*-containing nAChR, possibly a β4*-containing nAChR ^43,44^. Notably, the nicotinic EPSP is often followed by a mAChR-mediated IPSC/P responsible for a prolonged pause in THINs spontaneous activity. Although dual regulation of striatal FSIs by nicotinic and muscarinic recpetors has been shown, the two receptors were in distinct subcellular regions ^13^. However, functional modulation of postsynaptic membrane excitability in the same neuron via nAChR and mAChRs has not been shown in the striatum. While it is known that several populations of cortical and hippocampal interneurons express both nAChRs and mAChRs ^21,23^ their associated roles after the activation of a cholinergic input have not been described. However, mixed excitatory/inhibitory responses involving either mAChRs or nAChRs have been observed in several interneuron populations ^22^. Hence, we suggest that such precise regulation of GINs by cholinergic inputs exists in other systems, e.g., the striatum.

This mechanism of regulation of THINs activity by CINs should have a large influence on striatal functioning given the widespread targets of THINs (SPNs, LTSIs, CINs). Further, given the importance of other cholinergic systems in the generation of θ oscillation in the hippocampus, it is plausible that this oscillatory activity induced in several population of interneurons by CINs could be responsible for the propagation of θ oscillations in the striatum whose source(s) is still unidentified ^45,46^.

Another striatal GIN revealed in the Htr3a-Cre mice, the SABIs also receives suprathreshold nicotinic mediated excitation after optogenetic stimulation of CINs. SABIs have been described as a population of interneurons selectively targeting other interneurons ^7^ but not SPNs which could be compared to the VIP+ interneurons in the cortex. Comparably, VIP cortical interneurons express nAChRs and the resulting depolarization would cause a disinhibition of principal neurons essential for behavioral function ^47^. While the functional role of the CINs innervation of SABIs remains to be elucidated, it is likely that the large bursting activity induced by CINs optogenetic stimulation is sufficient to silence the striatal targets of SABIs resulting in the disinhibition of SPNs ^7^. Such disinhibition of SPNs has been suggested to play a role in important striatal function such as the transition from down state to upstate and/or the creation of cell assemblies ^48–52^.

Consistent with previous studies, we found that optogenetic stimulation of CINs evoke a long pause in the firing activity of LTSIs that is mediated mostly by mAChRs activation and a participation of GABA_A_ receptors certainly via CIN-activation of THINs ^5,6,14,53,54^. Here, in addition to confirming these data, we also show the presence of an EPSC/P in most recorded LTSIs that involve both nAChRs and glutamate receptor activation. While the nAChRs activation is consistent with previous reports demonstrating β2*-expressing nAChR expression by LTSIs ^14,44^, it is still unclear whether the glutamatergic response is due to the co-release of glutamate by CINs ^34^ or via presynaptic nAChR activation of glutamatergic afferents ^55–57^. Our pharmacological experiments favor the latter option as the presence of nAChR antagonists fully abolish the EPSC in most recorded LTSIs.

Finally, we showed that optogenetic activation of CINs triggers dual EPSC / IPSC in FSIs. Our results are consistent with previous reports suggesting the involvement of both glutamatergic and nAChRs in the EPSC ^13,34^ which may be due at least in part to the co-release of glutamate and ACh onto FSIs ^34^. Further, CINs optogenetic stimulation triggers GABA_Aslow_ in the majority of the recorded FSIs. The timing, the slow kinetic and the sensitivity to the blockade of GABA reuptake of the IPSC suggest the involvement of NGFs. However, while it has been suggested that FSIs innervate NGF interneurons ^51^, the reciprocal connection has never been reported. Remarkably, the blockade of the CINs-evoked GABA_Aslow_ with a GABA_A_ receptor antagonist revealed late glutamatergic asynchronous EPSC barrages suggesting the disinhibition and burst firing of striatal glutamatergic axons. Previous literature highlighted an important role of GABA in the regulation of corticostriatal glutamate release ^58–60^. In addition, GABAergic transmission can control the temporal window of spiking of SPNs by regulating the generation and propagation of plateau potentials, generated by clusters of excitatory inputs, essential for dendritic computation and spatiotemporal synaptic integration in SPNs ^60^. Interestingly, GABA_Aslow_ originating from NGF were the most efficient for dendritic plateau inhibition in SPNs ^60^. Consistently, the long-lasting EPSC barrages in FSI following blockade of GABA_Aslow_ were likely mediated by NGF interneurons which are heavily innervated by CINs ^2,44^. We propose that CINs activation of NGF interneurons tonically inhibits corticostriatal terminals that densely innervate FSIs ^25,61,62^, and that this disinhibition using a GABA_A_ antagonist is responsible for the asynchronous glutamatergic EPSCs measured in FSIs. This mechanism may be important for prolonging the period of membrane depolarization and input integration in FSIs.

When investigating the influence of THINs and LTSIs in the disynaptic inhibition of SPNs following CINs optogenetic activation, we found that LTS do not significantly participate, consistent with the modest excitation they receive from CINs. However, using double optogenetic stimulation we show that optogenetic inhibition of THINs provoke a significant decrease of the GABA_Afast_ and the GABA_Aslow_ in SPNs following activation of CINs. Our previous results demonstrate that THINs only evoke GABA_Afast_ in connected SPNs ^7,27,28^. Hence, the present data demonstrates that the reduction of the GABA_Afast_ current in SPNs following activation of CINs involves the intrinsic connectivity of THINs with CINs and SPNs. These results solidify our knowledge of the circuitry involved in the disynaptic inhibition of SPNs. Indeed, while it has been suggested that the GABA_Afast_ and the GABA_Aslow_ could originate from GABA released by dopaminergic afferents ^4^, our previous study demonstrated that GABAergic interneurons transduced in Htr3a-Cre mice (FSIs, NGFs, FAIs and SABIs) are involved in both components of the disynaptic inhibition of SPNs ^3^. However, the source of the GABA_Afast_ was still puzzling. Consistent with our present data, FSIs and FAIs do not seem to play a role ^3,4,10^. Given that SABIs do not innervate SPNs ^7^ they cannot participate in such circuit.

Here, we demonstrate that THINs are directly involved in the disynaptic GABA_Afast_ measured in SPNs following optogenetic stimulation of SPNs providing a mechanism through which THINs, under the influence of CINs, could rapidly regulate the activity of striatal outputs independently from any extrinsic cortical or thalamic input. The reduction of the GABA_Aslow_ following optogenetic inhibition of THINs was more surprising. Indeed, the CINs induced GABA_Aslow_ in SPNs has been definitively attributed to CINs activation of NGF interneurons, that are the source of GABA_Aslow_ in SPNs ^2^ which is most likely the result of GABA spillover acting on both synaptic and extrasynaptic GABA_A_ receptors ^63^.

We hypothesize that the reduction of the GABA_Aslow_ in SPNs after inhibiting THINs is indirect, through the concomitant hyperpolarization of NGFs. One possible mechanism would be the existence of heterotypic electrotonic coupling between NGFs and THINs. In this scenario, the hyperpolarization of THINs with HR3.0, would hyperpolarize NGFs which would result in the reduction of the disynaptic sIPSC. Here, we demonstrate the existence of such heterotypic electrical coupling, which had not yet been reported in the striatum. The existence of heterotypic gap junctions electrotonically linking NGF interneurons and other striatal GINs such as the THINs, suggest that striatal NGFIs are in position to control striatal network activity via electrical coupling and could contribute to oscillatory activity by synchronizing interneuron networks as observed in cortical and hippocampal circuits ^41,63,64^. In addition, it has been suggested that interactions between GABA_Afast_ generating interneurons and GABA_Aslow_ generating interneurons could also contribute to theta frequency modulation of gamma oscillation in other brain structures ^65–67^. Interestingly while theta oscillations have also been observed in the striatum (especially during locomotion) their source is still unknown. We suggest that striatal NGF interneurons could participate in the generation of striatal theta oscillations.

Overall, our data demonstrate a complex and heterogeneous mechanism of regulation of striatal GINs by CINs involving various subtypes of nAChRs, mAChRs, heterosynaptic glutamatergic and GABAergic receptors as well as the existence of functional heterotypic electrotonic coupling. These results give a central role to CINs, notably via the underestimated nAChRs, in striatal circuits and provide additional synaptic mechanisms elucidating their significant role in striatal related behaviors. Indeed, CINs have been demonstrated to play an important role in striatal related functions such as reward processing and cognitive flexibility ^12,68–70^ as well as in associated neurological disorders including Parkinson’s disease, obsessive compulsive disorders and Tourette. However, the precise physiological mechanisms responsible are yet to discover. The hypothesis that nAChRs, selectively expressed postsynaptically by interneurons in the striatum, may play an important part is tempting. Indeed, in the recent years several reports have demonstrated the importance of several populations of striatal GINs in similar functions ^5,71–73^. In this context, we propose that the role of CINs and GINs in striatal-related behaviors could be explained, at least partially, by the intricate circuitry presented here. Such study is thus necessary to answer these questions and lay out new organizational principles of the synaptic organization between striatal interneurons (as it has been outlined in other structures) where spontaneously active CINs would coordinate hierarchically in a sophisticated manner the activity of GINs thereby influencing in a semi-autonomous manner striatal output activity.

## Methods

### Animals

All procedures used in this study were performed in accord with the National Institutes of Health Guide to the Care and Use of Laboratory Animals and with the approval of the Rutgers University Institutional Animal Care and Use Committee.

Subjects were adult (3–6 months of age) ChAT-ChR2 mice (Tg(Chat-COP4*H134R/EYFP,Slc18a3) 6Gfng/J; Jackson Labs, Bar Harbor, MA, USA) crossed with either 1) BAC transgenic TH–Cre [Tg(TH–Cre)12Gsat; Gene Expression Nervous System Atlas (GENSAT) 2) Htr3a-Cre mice (Tg (HTR3a-Cre)NO152Gsat/Mmucd, University of Davis)^74^, or 3) SST-Cre mice (Sst-IRES-Cre, Stock No: 013044, The Jackson laboratory). Further, to test the presence of electrical coupling between THINs and NGF interneurons we used double transgenic TH–Cre crossed with (BAC) transgenic mice that express the humanized Renilla green fluorescent protein (hrGFP) (Stratagene) under the control of the mouse NPY promoter (NPY-GFP, stock 006417; The Jackson Laboratory). Double transgenic mice (ChAT-ChR2-EYFP::TH-Cre, ChAT-ChR2-EYFP::HT3Ra-Cre, ChAT-ChR2-EYFP::SST-Cre and TH-Cre::NPY-GFP) were genotyped and those found to be heterozygous positive for the transgene of interest were used for all recordings. Mice were housed in groups of up to four per cage and maintained on a 12-h light cycle with ad libitum access to food and water.

### Intracerebral viral injection

The surgery and viral injection took place inside a Biosafety Level-2 isolation hood. Mice were anesthetized with isofluorane (1-2.5%, delivered with O2, 1 L/min) and placed within a stereotaxic frame. Bupivacaine was injected under the scalp for local anesthesia at the site of the surgery. Coordinates to target the striatum were 0.6 mm anterior and 1.9 mm lateral to Bregma. A replication non competent adeno-associated virus (AAV5-CAG-Flex-tdTomato, University of North Carolina, Vector Core Services, Chapel Hill, NC or AAV5-EF1α-DIO-epNHR3.0-mCherry, Penncore or AAV5-EF1α-DIO-hChR2(H134R)-mCherry, University of North Carolina, Vector Core Services, Chapel Hill, NC) was delivered by glass pipette to two sites −2.5 mm and −3.0 mm ventral to brain surface for a total volume of 0.8 µL at 9.2 nl/ 5sec using a Nanoject II Auto-Nanoliter Injector (Drummond Scientific Company), after which the pipette was left in place for 10 minutes before being slowly retracted. Following viral injection, mice were treated with ketoprofen and buprenorphine for analgesia. Expression of the viral transgene was allowed for at least 4 weeks before animals were used for histology or physiology.

### Preparation of brain slices

Mice (3-6 months old) were deeply anesthetized with an intraperitoneal injection of 80 mg/kg ketamine and 20 mg/kg xylazine before being transcardially perfused with ice cold N-methyl D-glucamine (NMDG)-based solution containing (in mM): 103.0 NMDG, 2.5 KCl, 1.2 NaH2PO4, 30.0 NaHCO3, 20.0 HEPES, 25.0 Glucose, 101.0 HCl, 10.0 MgSO4, 2.0 Thiourea, 3.0 sodium pyruvate, 12.0 N-acetyl cysteine, 0.5 CaCl2 (saturated with 95% O2 and 5% CO2, pH 7.2-7.4). After decapitation, the brain was quickly removed into a beaker containing the ice-cold oxygenated NMDG-based solution before obtaining 300 µm parahorizontal slices using a vibratome (VT1200S; Leica Microsystems). Sections were immediately transferred to recover in well oxygenated NMDG-based solution at 35°C for 5 minutes, after which they were transferred to well-oxygenated normal Ringer’s solution at 25°C until placed in the recording chamber constantly perfused (2– 4 ml/min) with oxygenated Ringer’s solution at 32-34°C. Drugs were applied in the perfusion medium and were dissolved freshly in Ringer’s solution.

### Fluorescence and differential interference contrast imaging and recording

For recording striatal interneurons transduced with fluorescent AAV, slices were initially visualized under epifluorescence illumination with a high-sensitivity digital frame transfer camera (Cooke SensiCam) mounted on an Olympus BX50-WI epifluorescence microscope with a 40X long working distance water-immersion lens. Once a fluorescent interneuron was identified, visualization was switched to infrared–differential interference contrast microscopy for the patching of the neuron. Micropipettes for whole-cell recording were constructed from 1.2mm outer diameter borosilicate pipettes on a Narishige PP-83 vertical puller. The standard internal solution for whole-cell current-clamp recording of interneurons was as follows (in mM): 130 K-gluconate, 10 KCl, 2 MgCl2, 10 HEPES, 4 Na_2_ATP, 0.4 Na_2_GTP, pH 7.3. For recording the disynaptic inhibition of SPNs following optogenetic stimulation of CINs we used a CsCl-based internal solution containing the following (in mM): 125 CsCl, 0.1 EGTA, 10 HEPES, 2 MgCl2, 4 Na2ATP, and 0.4 Na2GTP. This solution also contained 0.2% Alexa Fluor 594 (Molecular Device), or biocytin (Sigma) to verify the identity of SPNs and interneurons. Pipettes had a DC impedance of 3-5 MΩ. Membrane currents and potentials were recorded using an Axoclamp 700B amplifier (Molecular Devices). Recordings were digitized at 20 kHz with a CED Micro 1401 Mk II and a PC running Signal, version 5 (Cambridge Electronic Design). Optogenetic stimulation was performed using a high power Multi LED (LZC-R0H100-0000, 730 nm, 665 nm, 525 nm, 450 nm, 400 nm, 590 nm, mouser). Stimulation of channelrhodopsin-2 (ChR2)-expressing neurons in vitro consisted of 2 ms duration blue light pulses (450 nm). Optogenetic stimulation of HR3.0 consisted of 700 ms duration yellow light pulses (590 nm) starting 200 ms before the blue light pulse. Optogenetic pulses were delivered at 30 s intervals.

### Analyses

*Disynaptic inhibition of SPNs.* Because we hypothesized that the fast and slow components of the CIN-evoked IPSC involve different populations of striatal interneurons ^2,3^, we aimed to characterize these two components independently in order to determine their selective reduction after the inhibition of the different subtypes of interneurons tested transduced with halorhodopsin. Although the peaks of the two IPSC components constitute a variable mixed response of GABA_Afast_ and GABA_Aslow_, they represent their respective maximal contributions to the response and occur with a substantial delay between them. Therefore, we measured the current reduction at both time points.

### Drugs

Drugs that were used include bicuculline (10μM, Sigma) to block GABA_A_ receptors, Dihydro-β-erythroidine hydrobromide (DhβE; 1μM, Tocris) to block type 2 nicotinic receptors (containing β2-subunits), methyllycaconitine citrate (MLA; 500nM, Tocris) as an antagonist of type I nicotinic receptors (containing α7-subunits), mecamylamine hydrochloride (MEC; 5 μm, Tocris), Atropine (Sigma, 10μM) to block muscarinic receptors, CNQX (10μM, Tocris) and APV (10μM, Tocris) to block respectively AMPA and NMDA glutamate receptors. To block gap junction (electrotonic) communication we used carbenoxolone (100μM, Tocris).

### Imaging

Mice were deeply anesthetized with an intraperitoneal injection of 80 mg/kg ketamine and 20 mg/kg xylazine. Brain tissue was fixed by transcardial perfusion of 10 mL of ice-cold artificial cerebrospinal fluid (adjusted to 7.2-7.4 pH), followed by perfusion of 90-100 mL of 4% paraformaldehyde, 15% picric acid diluted in phosphate buffer. Fixed brains were extracted and post-fixed overnight in the same fixative solution. 50 μm sections were cut on a Vibratome 3000. Sections were mounted in Vectashield (Vector Labs, Burlingame, CA) and representative photomicrographs were taken using a confocal microscope (Fluoview FV1000, Olympus). Photomicrographs were taken using 10 and 40x objectives. Comparable pictures were taken using the same laser settings.

After whole-cell recording, slices containing biocytin-filled neurons were transferred into 4% paraformaldehyde with 15% picric acid in 0.1 M PB for overnight fixation (4°C). After multiple washes in PBS, sections were incubated in PBS containing 0.05% Triton (3h) followed by a blocking solution containing normal donkey serum (10%) and 0.05% Triton (2h). Then, sections were incubated overnight with Streptavidin, Alexa Fluor 405 conjugate (Thermofisher, 1:1000) and antibody against GFP (rabbit polyclonal anti-GFP, Invitrogen, 1:1000). Sections were then mounted in a fluorescence medium (ProLong Glass Antifade Mountant, Invitrogen) and representative photomicrographs were taken at 10X and 40X using a confocal microscope (Fluoview FV1000, Olympus). Images were analyzed using ImageJ software.

### Statistical Analyses

Data are represented as mean ± SEM unless indicated otherwise. Comparisons were made using two-tailed paired Student’s t-test or one-way ANOVA followed by Tukey’s post hoc test, when appropriate. The statistical test used as well as numbers (n), p values and degree of freedom are indicated in the respective results section. Data are considered significant if p<0.05. Correlations between reduction of GABA_Afast_ and GABA_Aslow_ were done using a Pearson r correlation allowing the performance of a linear correlation between these two data sets.

## Data Availability

The authors declare that the data supporting the findings of this study are available within the main manuscript and the supplementary information files. However, additional supporting data are available from the corresponding author upon reasonable request.

## Acknowledgements

We would like to thank Shubhi Yadav and Mina Nakhla for excellent technical assistance. This research was supported by a NINDS R01 NS034865 to J.M.T., a NARSAD Young Investigator grant from the Brain and Behavior Research Foundation to M.A. and a Rutgers Busch Biomedical Grant to M.A. as well as Rutgers University.

## Author contributions

M.A. and J.M.T. designed the study. S.K., M.A. and B.G. collected the electrophysiology data. F.S. participated in collecting anatomical data and managed the transgenic colony. M.A. wrote the first draft of the manuscript with major inputs from S.K. and J.M.T.

## Supplementary Information

**Supplementary Figure 1.**
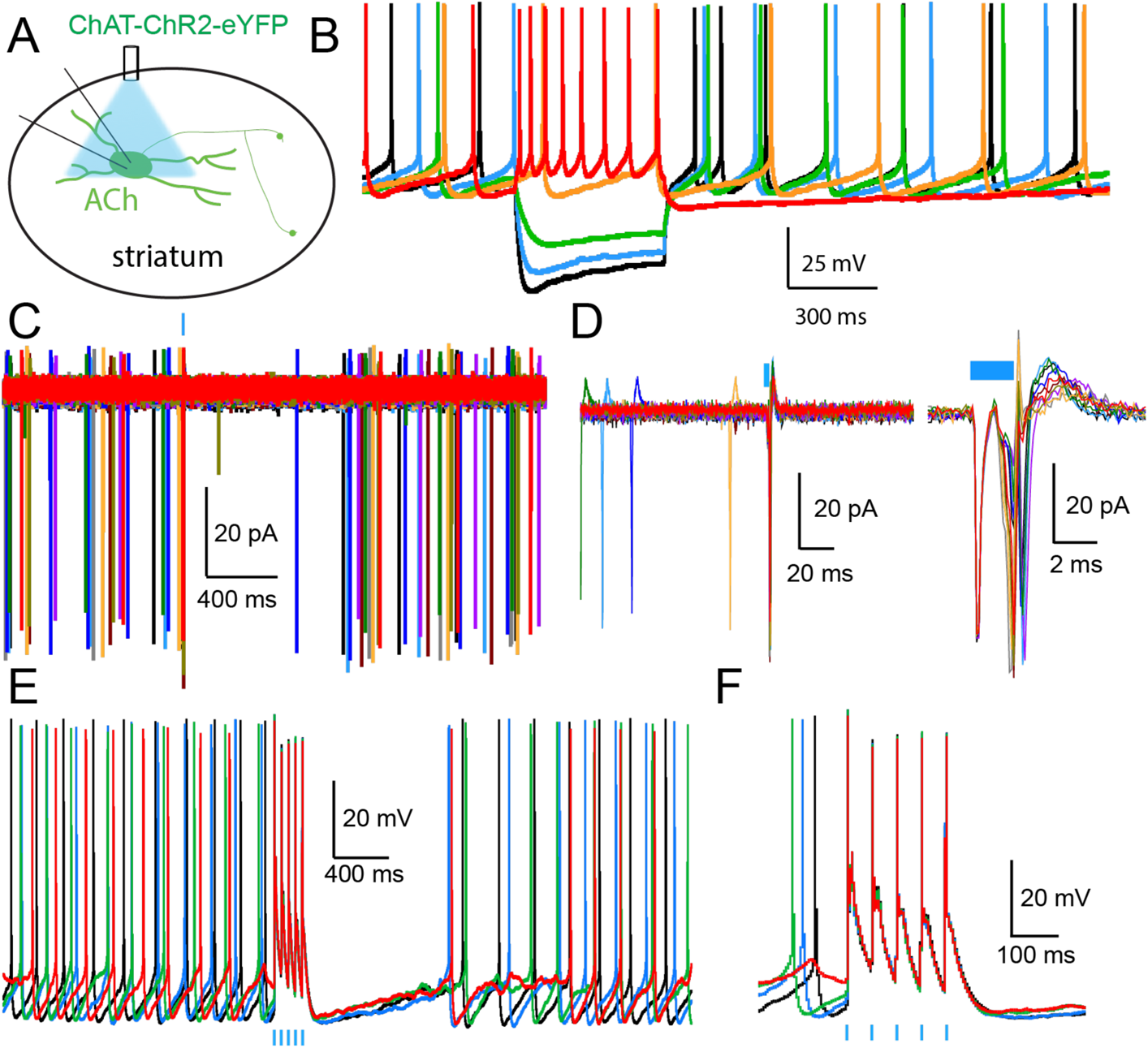
Optogenetic stimulation of ChR2-expressing CINs. **A**. Schematic describing the experimental design recording CIN in ChAT-ChR2-eYFP transgenic mice. **B**. Typical response of a CIN to somatic current injection. Note the presence of I_h_ sag current and the spontaneous activity. **C, D.** Cell attached recording of a ChR2-expressing CIN responding to a blue light pulse (2ms) with an AP. **E-F.** Current clamp recording of a ChR2-expressing CIN responding to a train of blue light pulses (5 pulses, 20Hz) with APs.

**Supplementary Figure 2.**
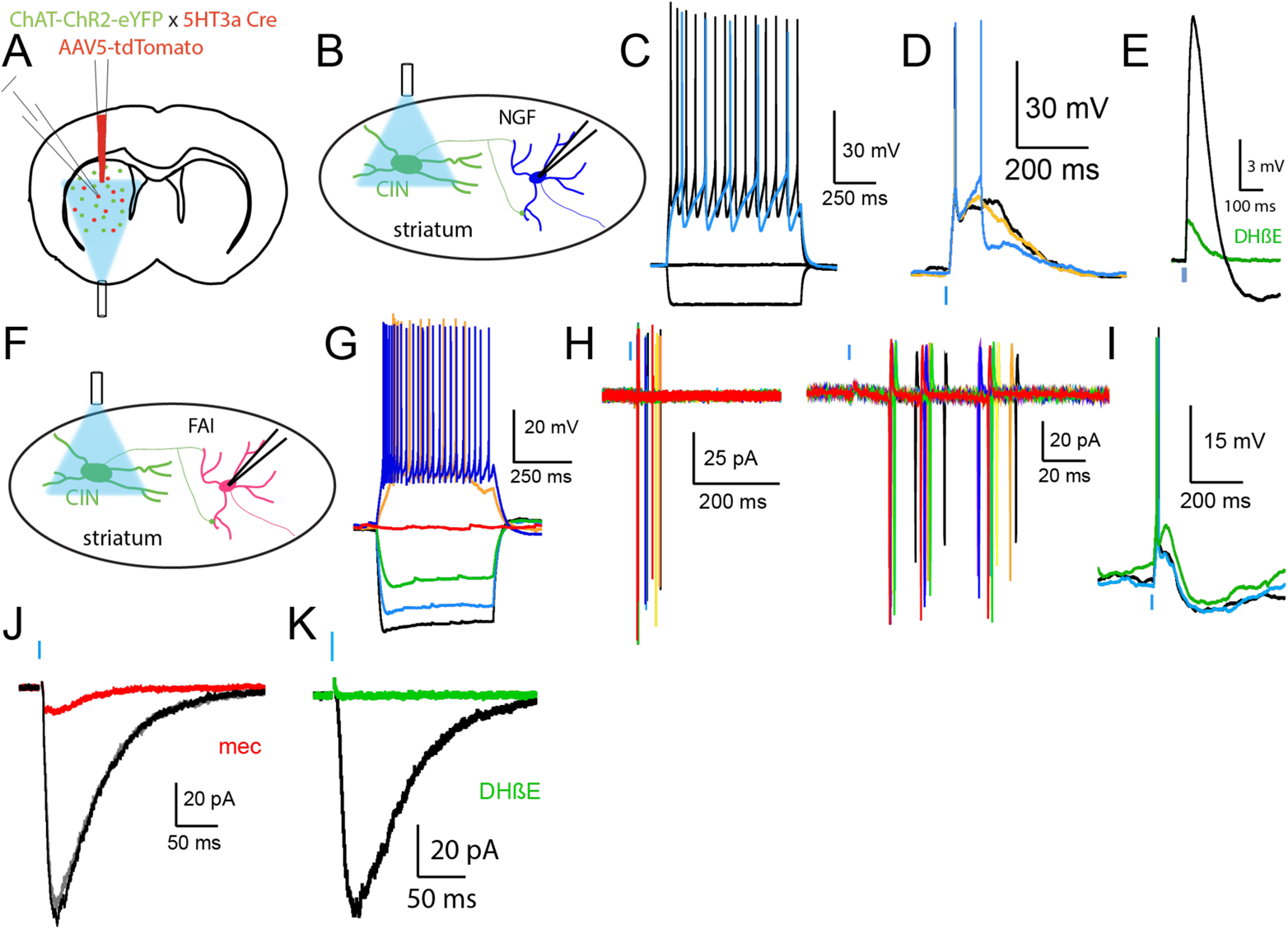
Responses of NGFs (B-E) and FAIs (F-K) to optogenetic stimulation of CINs. **A.** Schematic describing the experimental design using double transgenic ChAT-ChR2::Htr3a-Cre mice injected with a Flex-tdTomato AAV in the striatum. **B.** Recording of tdTomato transduced NGF responses to optogenetic stimulation of CINs. **C.** Typical responses of NGF interneuron to somatic current injection. **D.** Stimulation of CINs (blue bar) elicit large depolarization and AP firing in a NGF. **E.** The EPSP can be almost abolished by bath application of DHβE (1μM, see ^2^). **F.** Recording of tdTomato transduced FAIs responses to optogenetic stimulation of CINs. **G.** Typical responses of a FAI interneuron to somatic current injection. **H-I** Stimulation of CINs evokes AP firing in FAIs in cell attach (**H**) and current clamp (**I**). The CIN-induced EPSC in FAI can be block by MEC (5μΜ) or DHβE (1μM, see ^10^).

**Supplementary Figure 3.**
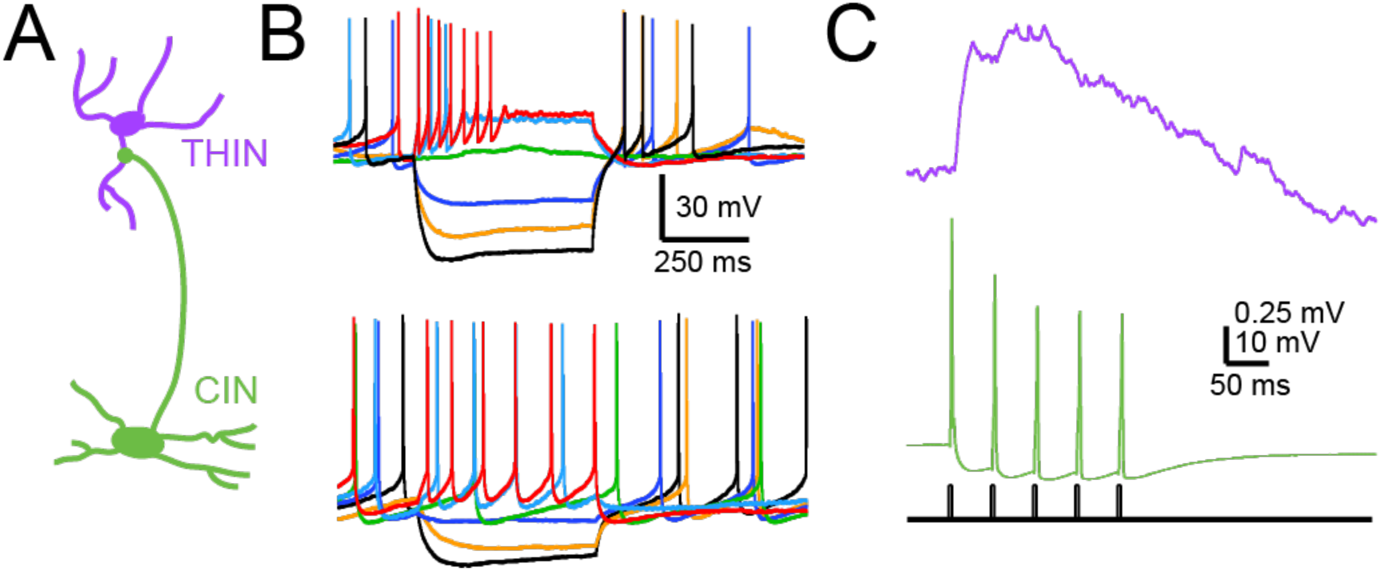
Paired-recording between CINs and THINs. **A.** Schematic of the recording design **B.** Typical responses of a THIN (top) and CIN (bottom) to somatic current injection. **C.** A train of 5 APs elicited in a CIN (bottom, green) induce an EPSP in a synaptically connected THIN (top, purple, see ^1^)

**Supplementary Figure 4.**
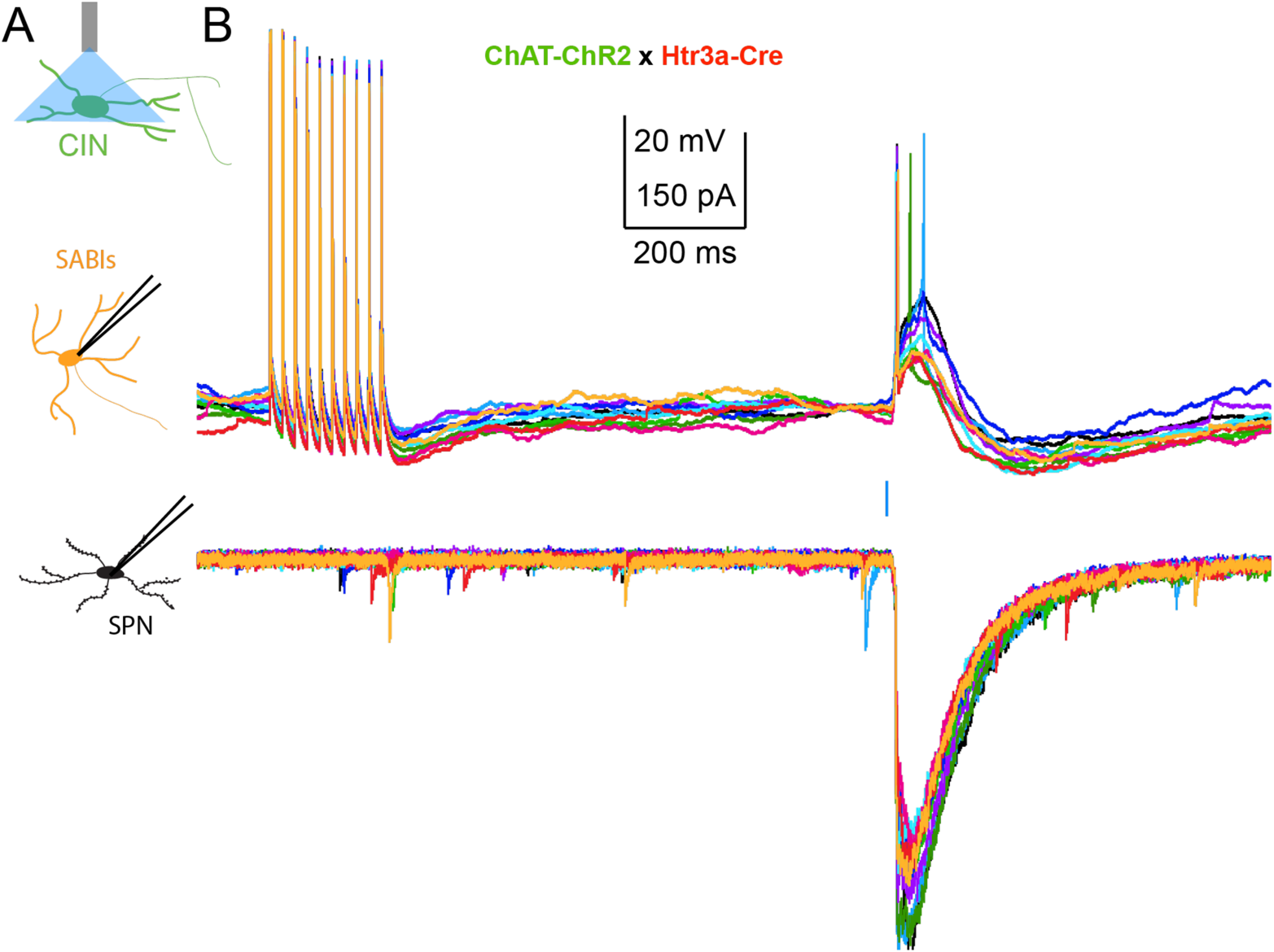
Paired-recording between a SABI and a SPN combined with CINs optogenetic stimulation. **A.** Schematic demonstrating the experimental design where we recorded from a pair of SABI and SPNs and measured their connectivity as well as synaptic response to optogenetic stimulation of CINs. We used double transgenic ChATChR2-eYFP::Htr3a-Cre mice injected with a Flex-tdTomato AAV. **B.** Top: We elicited a train of APs in a SABI recorded in current clamp (10 pulses, 20Hz). Bottom: The SPN recorded in voltage clamp (CsCl Internal solution, see methods) is not synaptically connected with the SABI (see ^7^). CINs optogenetic stimulation (2 ms, blue bar) evoke large depolarization and AP firing in the SABI as well as large disynaptic IPSC in the SPN.

**Supplementary Figure 5.**
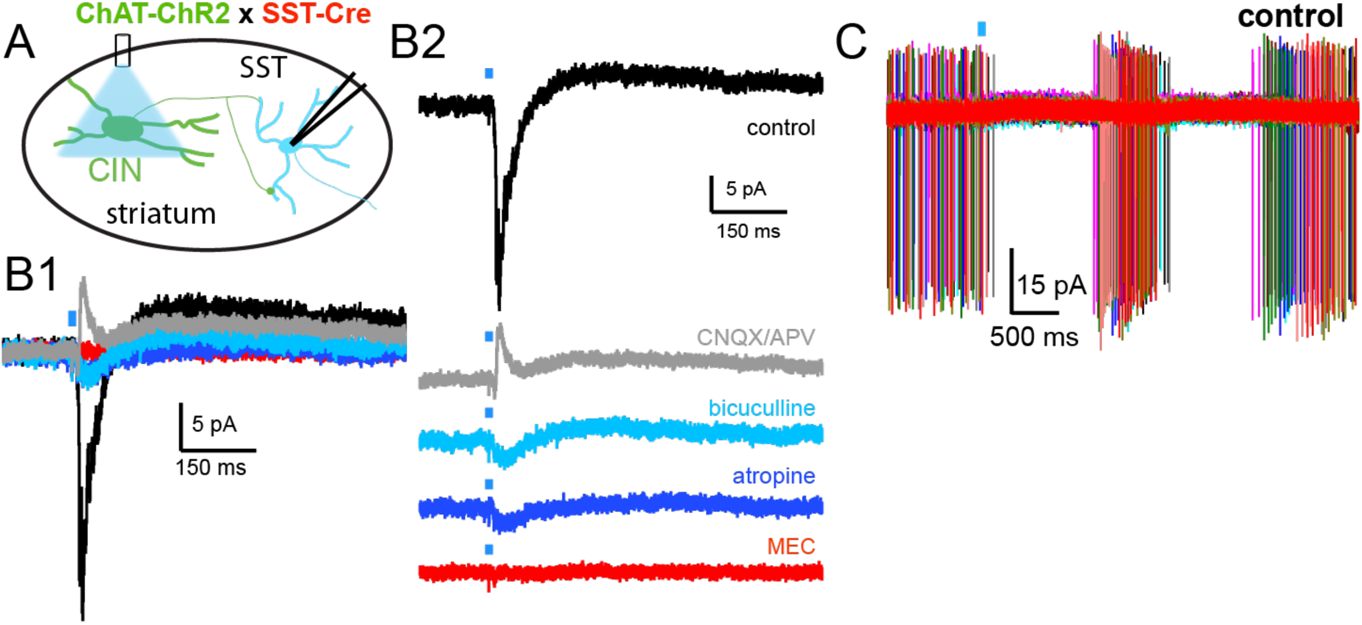
Cholinergic synaptic connectivity with LTSIs. **A.** We used double transgenic ChAT-ChR2-eYFP::SST-Cre mice injected with a Flex-tdTomato AAV and recorded from transduced LTSIs. **B.** Optogenetic stimulation of CINs evokes a fast EPSC followed by a slow IPSC (black). Application of glutamate receptor antagonists (cnqx, 10μΜ and APV, 10μΜ) revealed a fast disynaptic IPSC in n=4/8 LTSI. This IPSC can be blocked by a GABA_A_ receptor antagonist (bicuculline, 10μM), suggesting the action of a disynaptic circuit involving CIN-THINs-LTSIs (see ^5,6^). Further, the slow IPSC involve muscarinic receptor activation (Atropine, 10μM) and the small remaining EPSC can be blocked by a nicotinic receptor antagonist (MEC, 5μM). **C.** In some instance (∼25%), LTSIs recorded in cell attach respond to CINs optogenetic stimulation with burst firing activity ^6^.

**Supplementary Figure 6.**
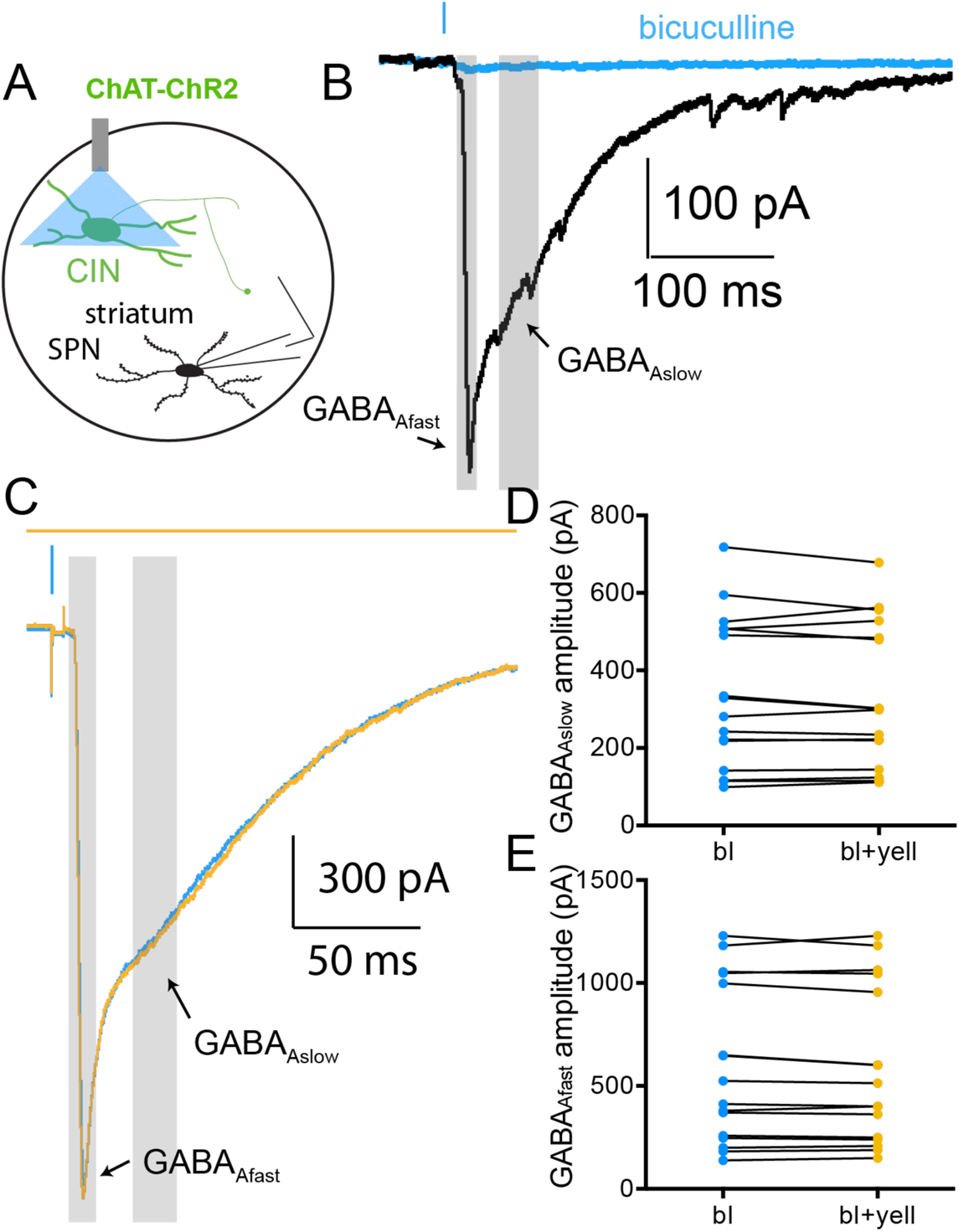
Light control for the nicotinic-mediated disynaptic inhibition of SPNs. **A**. Schematic illustrating the experimental design. SPNs were recorded in ChAT-ChR2-eYFP mice. **B**. Optogenetic stimulation of CINs evokes large disynaptic IPSCs composed of a fast and slow IPSC (GABA_Afast_ and GABA_Aslow_, ^2–4^). **C**. Yellow light illumination (700 ms starting 200ms before blue light pulse) does not significantly affect the GABA_Afast_ (quantified in **E**) or the GABA_Aslow_ IPSC (quantified in **D**).

## Notes

### Competing Interest Statement

The authors have declared no competing interest.

